# Design principles underlying nearly-homeostatic biological networks

**DOI:** 10.1101/2025.10.23.683559

**Authors:** Zhe F. Tang, David R. McMillen

## Abstract

A nearly-homeostatic system like body temperature maintenance keeps the steady state system output like internal body temperature within a narrow range, regardless of different persistent levels of environmental perturbations like external temperatures. Nearly-homeostatic systems can be implemented to guarantee performance of a therapeutic device in different patient contexts. Exploration of different near-homeostasis-supporting architectures to satisfy different design requirements is necessary, but methods for doing so in a fast and comprehensive manner remain elusive. We have identified two constrained optimization approaches to find near-homeostasis supporting architectures 10 to 100 times faster than brute-force search. Once such architectures are found, characterizing the underlying mechanisms of near-homeostasis is hindered by the very cumbersome and limited nature of traditional statistical analysis approaches. We have developed two levels of “inverse homeostasis plots”, and used them to identify two novel near-homeostasis mechanisms that tolerate much higher undesired basal expression and much higher undesired first order degradation rates than previously characterized near-homeostasis mechanisms.

**Teaser:** A comprehensive toolset was developed for much faster discovery and much more comprehensive analysis of near-homeostasis-supporting architectures.

## Introduction

Homeostasis is a prevalent concept in biology. A homeostatic system like internal body temperature maintenance keeps the steady state system output like internal body temperature at a fixed set point, despite different persistent levels of environmental perturbations like external temperatures (*1,2*) (Fig. 1A). A nearly homeostatic system maintains the steady state output within some narrow range, regardless of different persistent levels of environmental perturbation (Fig. 1A).

**Fig. 1:**
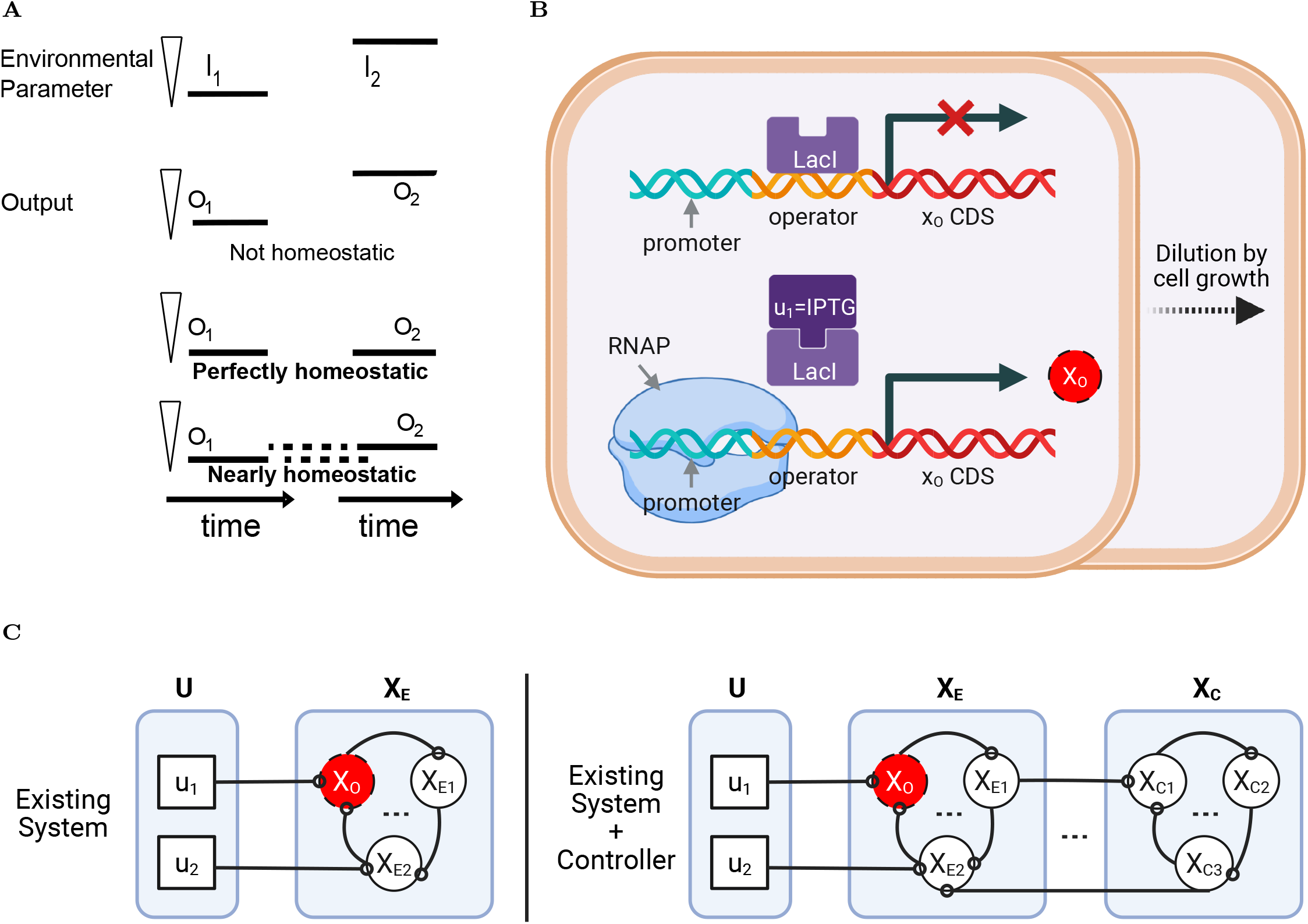
An introduction to homeostatic systems. (A) Definition of homeostasis and near-homeostasis. (B) A basic transcriptional element to be modeled. **??** A homeostatic system can ensure therapeutic molecular level (green dots) is independent of nutrient state (squiggly fragments) or microbiotic competitive landscape. (C) Definition of implementing a system to ensure near-homeostasis of the output *x*_*O*_’s steady state.

It would be useful to be able to construct synthetic nearly-homeostatic systems for various applications in biological engineering and medicine. For example, one could engineer bacteria to secret therapeutic molecules (shown in green) whose concentration depends on a sensed level of gut inflammation, while resisting perturbations applied by patient-to-patient variations such as nutrient states and the influence of the native gut flora. Such perturbations could otherwise greatly perturb therapeutic molecule levels.

Our central goal is to identify commonly occurring qualitative regulatory patterns and quantitative regulatory strengths that support near-homeostasis. More precisely, we start with an existing system ***x***_***E***_ = (*x*_*O*_, *x*_*E*1_, *x*_*E*2_, …) that is affected by a set of perturbations ***u*** = (*u*_1_, *u*_2_), and seek a set of controller components ***x***_***C***_ = (*x*_*C*1_, *x*_*C*2_, *x*_*C*3_, …) that keeps the steady state output *x*_*O*_ within a predefined narrow range despite different persistent perturbation levels (Fig. 1C). The qualitative regulatory patterns of how the system components are activated or repressed by each other is called topology of the system. Each activation or repression is further characterized by a set of unique regulatory strengths. A topology and a class of regulatory strengths associated with that topology is called a network architecture.

Different architectures of nearly-homeostatic systems may be suitable for different purposes, so architecture exploration is a necessary step prior to the implementation of a nearly-homeostatic system. In the engineered bacteria example, it would be desirable to minimize magnitude and duration of oscillations in the therapeutic molecule level as the microbial environment in the gut changes. On the other hand, when it comes to maintaining blood glucose steady state by an insulin pump, it becomes crucial to avoid even a transient amount of extra acting insulin in the body that dramatically increases the risk of acutely dangerous hypoglycemia (*3, 4*). Because it takes time for subcutaenously injected insulin to move to the bloodstream to start decreasing blood glucose levels and not all patients have identical insulin movement and clearance rates, it would be useful to explore a variety of controller architectures to find one that minimize the amount of extra insulin in the bloodstream.

Previously identified near-homeostasis supporting architectures have limited applicability in new therapeutic settings, making rational design alone insufficient for architecture exploration. For instance, although negative feedback networks and incoherent feedforward loops were the only types of topologies among metabolic networks (*1*) and transcriptional networks (*5*) that were capable of near-homeostasis in previous studies, only a small fraction of the possible regulatory strengths within each correct topology actually supported near-homeostasis and the stated classes of regulatory strengths for the same topology were different between the two network types. This lack of generalization capability for near-homeostasis supporting architectures is even true for the famous antithetic integral controller (*6, 7*), where the unavoidable dilution of the controller components compromises the homeostatic performance (*7*). Thus, it would be useful to have a fast and generalizable toolkit that finds and characterizes the majority of the possible near-homeostasis supporting architectures for any given experimental system.

In this study, we have explored a variety of different approaches to find near-homeostasis supporting architectures that are faster than brute force search while being equally comprehensive; once enough near-homeostasis supporting architectures were found, we then built a framework to identify the underlying mechanisms of near-homeostasis much more readily than the traditional trial-and-error based statistical analysis. We first explored the feasibility of many previously published approaches to find near-homeostasis supporting architectures in terms both speed and comprehensiveness compared to brute force search (*8*,1,2,9,10). Taking into account inherent limitations of prior approaches, we characterized four numerical optimization methods to find near-homeostasis supporting architectures. We found two constrained optimization approaches that were able to find near-homeostasis supporting architectures 10 to 100 times faster than the brute-force search.

The optimization approaches are fast for both high-dimensional elementary chemical reaction networks and higher-level Hill function based transcriptional networks. We then proceeded to identify the mechanisms responsible for near-homeostasis, and characterize the advantages of the discovered mechanisms. Since the prior approach of statistical analysis to identify mechanisms of near-homeostasis was quite cumbersome and not comprehensive, we developed a novel method (using two levels of what we denote as “inverse homeostasis plots”) to easily identify over 80% of near-homeostasis mechanisms for both transcriptional networks and arbitrary chemical reaction networks. By combining mechanism identification with rational design, we discovered two novel near-homeostasis mechanisms that tolerate much higher undesired basal expression and much higher undesired first order degradation than previously characterized near-homeostasis mechanisms.

## Results

### Mathematical preliminaries

First order Ordinary Differential Equations (ODEs) are useful descriptors of nearly-homeostatic biological systems: a set of state variables represents the number or concentration of a biological molecule or complex, and each state variable has a rate of change, specified in terms of the current state of the system. Suppose our existing system *A*_*E*_ is described by a set of first order ODEs describing the rate of change of the concentration of each biological component of the system in a growing cell

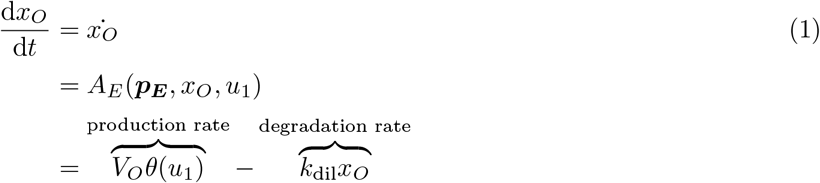

In our example, perturbation *u*_1_ = IPTG activates transcription of the scalar system output *x*_*O*_, by deactivating LacI repressor 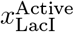 bound to the promoter of the *x*_*O*_ coding sequence (Fig. 1B). This first order ODE essentially says that the current rate of change in the system output *x*_*O*_ depends on a fixed set of regulatory parameters ***p***_***E***_, the current level of the system output itself *x*_*O*_, and a fixed perturbation level *u*_1_. The rate of change of the system output is its production rate subtracted by its degradation rate. Here, the production rate of the system output is simply the maximal production rate *V*_*O*_ multiplied by a fraction of available promoters that depends on the perturbation *u*_1_ = IPTG, *θ*(*u*_1_). The degradation rate of the system output is assumed to be dominated by first-order dilution of the system output due to cell growth. The maximal production rate *V*_*O*_, cell dilution constant *k*_dil_, and any constants within the function *θ*(*u*_1_) are all coordinates of the regulatory parameter vector ***p***_***E***_. At steady state, the rates of all system components equal zero

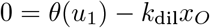

We can solve for the steady state of the system output at each perturbation level *u*

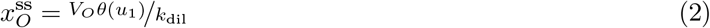

The relative sensitivity function is the rate of change in the steady state output with respect to the change in perturbation *u*_1_ further divided by the level of steady state output:.

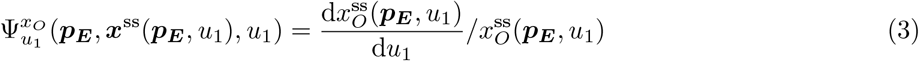

A system is nearly-homeostatic at *u*_1_ if the magnitude of the relative sensitivity at *u*_1_ is less than the desired threshold defined by the design requirements of a particular application. If there are multiple perturbations to the system ***u*** = [*u*_1_, *u*_2_, …, *u*_*N*_], a system is empirically nearly-homeostatic if the magnitude of empirical relative sensitivity at the first and each of the other perturbation vectors {***u***_**1**_, ***u***_**2**_, …, ***u***_***N***_ }

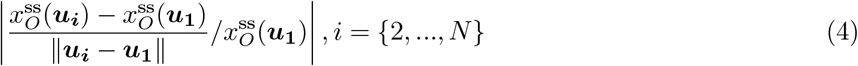

is less than the desired threshold.

Here, we restate our central goal in the context of mathematical models for regulatory networks. To make *x*_*O*_ of the existing system ***x***_***E***_ nearly-homeostatic, we want to find a controller architecture composed of how the controller works internally ***A***_***C***_, how the controller acts upon the existing system ***A***_***CE***_

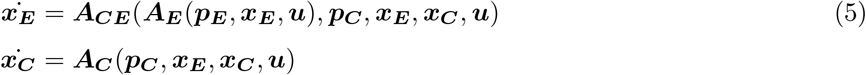

such that whenever a persistent perturbation ***u*** = (*u*_1_, *u*_2_) is within the desired range 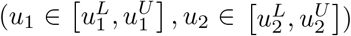, steady state output 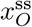 is within the desired range 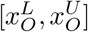 as well (Fig. 1C). The controller architecture encompasses the qualitative description of how the system components (***x***_***E***_, ***x***_***C***_) are activated or repressed by each other, which is also called the topology of the system. For example, the topological parameter describing the regulation of *x*_*C*1_ by *x*_*C*2_ can take on three discrete values, corresponding to repression, no regulation, and activation. The second component of the controller architecture consists of a class of non-topological parameter sets. Non-topological parameters are the regulatory strengths of different interactions within the network, and also are the complement of topological parameters within the regulatory parameters (***p***_***E***_, ***p***_***C***_). Before identifying a specific mechanism of near-homeostasis, one can arbitrarily define a class of non-topological parameter sets by selecting any region of non-topological parameter values to be associated with the network architecture. If a mechanism of near-homeostasis is well defined, one can associate the network architecture only with the non-topological parameter sets which all have that same underlying mechanism.

### No prior approach to find near-homeostasis supporting architectures is both faster and as comprehensive compared to brute-force search

There is currently no approach to find near-homeostasis supporting architectures that is both comprehensive and faster than brute-force search. In a comprehensive brute force approach, Ma et al (*1*) defined fixed topologies and randomly drew thousands of non-topological parameter sets for each of them, found a steady state at each of two fixed perturbations, and determined whether the empirical relative sensitivity was sufficiently small (Fig. 2A). For a three component metabolic network where one system component can be down regulated, not regulated, or up-regulated by another system component, there are already 3^3×3^ = 19, 683 possible topologies. The comprehensive search strategy quickly becomes intractable beyond three system components because each additional system component exponentially increases the number of possible topologies. However, if one only cares about finding a number of near-homeostasis parameter sets without restriction to a particular topology, one obvious method to speed up the comprehensive brute force search is to randomly hop between different topologies and non-topological parameter sets (Fig. 2B). For systems with more than two perturbation levels, another obvious speed-up method is to stop finding steady states upon encountering the first perturbation level that does not permit empirical near-homeostasis. This faster approach will serve as a baseline to compare with other approaches to find near-homeostasis supporting architectures.

**Fig. 2:**
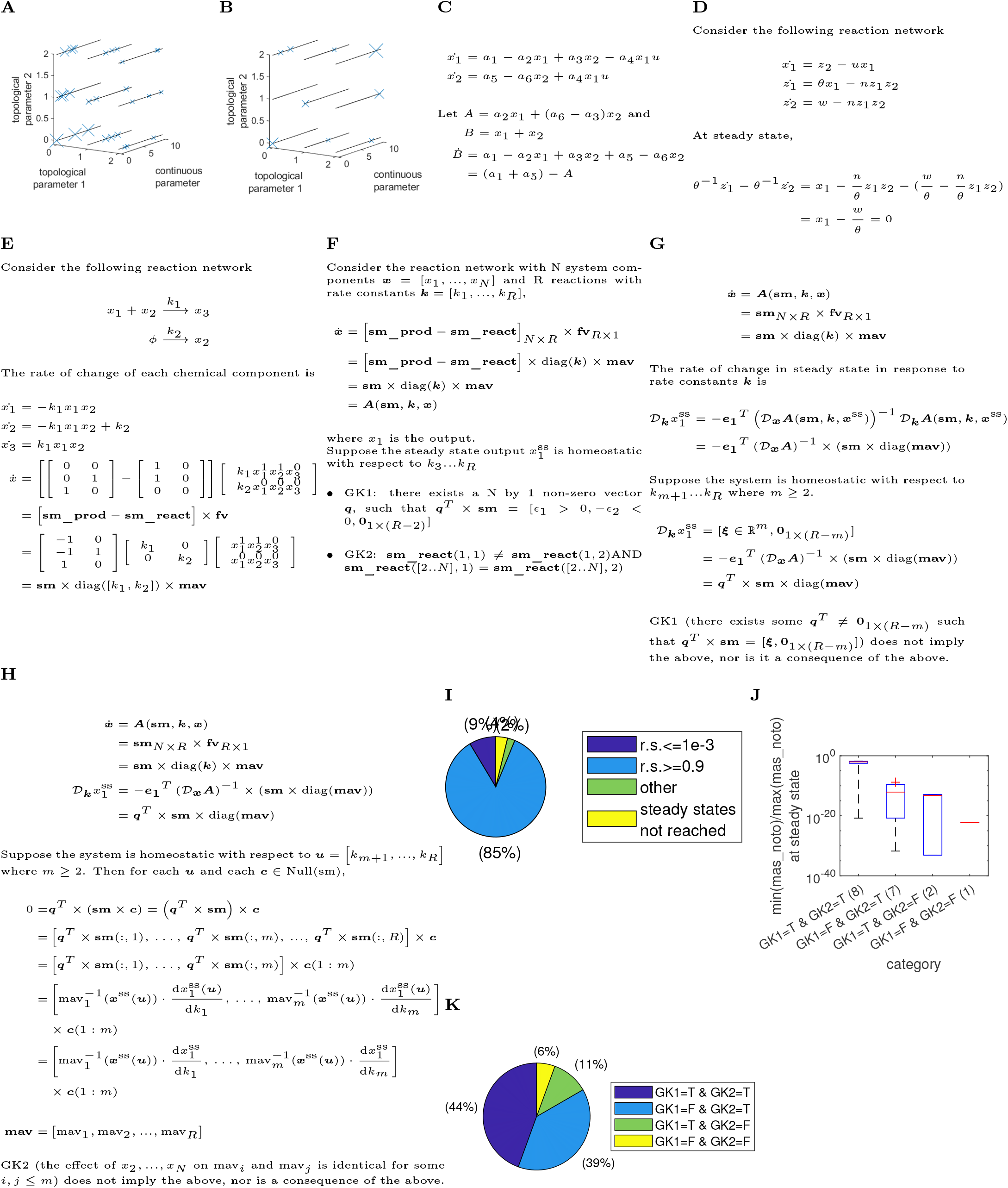
Analysis of prior approaches to find near-homeostasis supporting architectures. (A) Schematic of the comprehensive brute-force search (*1*). The decreasing size of data points corresponds to the increasing order in which the search occurs. (B) Schematic of the optimized brute-force search. The decreasing size of data points corresponds to the increasing order in which the search occurs. (C) Analyzing a perfectly homeostatic system using the internal model principle (*8*). (D) Analyzing a perfectly homeostatic system using elimination polynomials (*10*). (E) Describing a simple reaction network using stoichiometric matrices. (F) Gupta et al (*9*)’s conditions (GK1, GK2) for perfect homeoestasis. (G) Condition GK1 is not necessary or sufficient when the set point is encoded by 3 or more reactions. (H) Condition GK2 is not necessary when the set point is encoded by 3 or more reactions. (I) Relative sensitivity (r.s.) of parameter sets satisfying both GK1 and GK2. (J) Characterization of nearly-perfect-homeostasis supporting parameter sets based on Gupta et al (*9*)’s conditions and how close the smallest non-set-point-encoding mass action is to zero. A parameter set is nearly-perfectly homeostatic if its relative sensitivity is less than 5 × 10^−5^. Because simulated systems often arrive near theoretical steady states rather than arrive at the exact theoretical steady state, perfect homeostasis supporting parameter sets often appear as supporting nearly-perfect-homeostasis. “mas_noto” means mass actions that does NOT encode the set point of the output 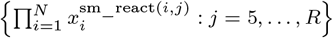. (K) Proportion of nearly-perfect-homeostasis supporting parameter sets satisfying Gupta et al (*9*)’s conditions.

Although perfectly homeostatic systems have been frequently analyzed using the internal model principle (IMP) of control theory, we have not found the IMP to be sufficiently predictive of parameter conditions for perfect-homeostasis. The internal model principle (IMP) states that if a nonlinear system is perfectly-homeostatic, then there exists a nonlinear transformation of system components such that the solutions of a subsystem after the nonlinear transformation can be mapped to the set of persistent perturbations (*8*). Suitable nonlinear transformations have been found for already-perfectly-homeostatic systems to demonstrate satisfaction of the internal model principle (*8*). In their example (Fig. 2C), the solutions of transformed subsystem *B* are a range of constant values that can be identically mapped to the set of persistent perturbations. However, we and others (*10*) do not believe there is a straightforward algorithm to tell us under what parameter conditions the IMP would be satisfied.

Araujo et al (*10*) were able to compute polynomials to eliminate elements in the rate equations in a way that makes the system output separable in perfectly homeostatic systems. Suppose we have the classic antithetic integral controller. We can compute an elimination polynomial coefficient for each rate equation such that the sum of the polynomial weighted rate equations results in a separable output term (Fig. 2D). For more complicated systems, the elimination polynomials were computed using the nonpolynomial-time Grobner basis algorithm. Since the Grobner basis algorithm is only guaranteed to complete within polynomial time for perfectly homeostatic systems but can have non-polynomial run-time for not homeostatic systems (*10, 11*), Araujo et al (*10*)’s approach would unlikely find perfect-homeostasis supporting architectures faster than brute force search. Other algebraic methods (*2, 12*), including one from our own group (*2*), also suffer from the issue of high computational complexity. Those methods required analyzing exponential numbers of algebraic terms in the sensitivity function 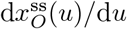 as the number of system components increase, making the methods unlikely to be faster than brute-force search.

Lastly, Gupta et al (*9*) devised two very fast and simple algebraic tests to determine whether an elementary reaction network is perfectly homeostatic. Elementary reaction networks are an important class, since any biological regulatory network can in principle be represented as an elementary reaction network by including a sufficiently complete set of biochemical reactions. Rate equations for reaction networks can be organized neatly into the product of matrices, making such systems suitable for application of standard linear algebra tools. Here, suppose we have two reactions where *x*_1_ and *x*_2_ combine together to form *x*_3_, and *x*_2_ is constantly being produced at a fixed rate (Fig. 2E). We can easily write law of mass action based rates of change for each of *x*_1_, *x*_2_, and *x*_3_. All reactant coefficients can be organized into the reactant stoichoimetric matrix. All product coefficients can be organized into the product stoichoimetric matrix. The rates of all reactions can be organized into the flux vector. Therefore, the rates of this reaction network can be stated in terms of the reactant stoichiometric matrix (sm_react), product stoichoimetric matrix (sm_prod), and the flux vector (fv). Each flux vector can be decomposed into the rate constants themselves and the mass action vector (mas). The two algebraic tests (GK1, GK2) essentially test for suitability of product and reactant stoichiometric coefficients for perfect-homeostasis (Fig. 2F). Although GK1 and GK2 are very quick to check, the conditions are restricted to finding reaction networks where the homeostatic set point is encoded by only two rate constants.

We wanted to see if the two algebraic tests could be modified to be valid when the set point was encoded by more than two rate constants, and discovered inherent limitations that makes it difficult to generalize the two tests. We vastly simplified the proofs of GK1 and GK2 to focus on the underlying intuition. To derive GK1 (Fig. 2G), the original authors computed the sensitivity of the steady state output to every rate constant 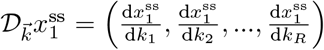. Homeostasis of the steady state output with respect to all rate constants except the first *m* set-point encoding rate constants means that the steady state sensitivity vector takes on the following form:

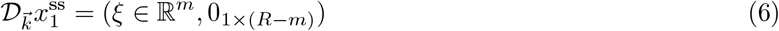

However, GK1 does not imply the existence of a system steady state that allows the sensitivity vector to attain the above form. This lack of a steady state causes many parameter sets satisfying GK1 and GK2 to fail. For example, in a four component reaction network with seven reactions and two set point encoding reactions, the number of parameter sets whose steady state output is far-from-homeostatic (Fig. 2I: r.s. > 0.9) is at least 10 times larger than the number of near-homeostasis supporting parameter sets (Fig. 2I: r.s. < 1 × 10^−3^). Additionally, a homeostatic system with a zero non-set-point encoding mass of action element among {mav_*m*+1_, …, mav_*R*_} does not necessarily permit a non-zero vector *q* that satisfies GK1, contradicting the idea that GK1 is a necessary condition for near-homeostasis. Consistently, the distribution of the relative minimum of non-set-point encoding mass of action elements {mav_*m*+1_, …, mav_*R*_}

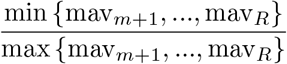

is much lower when GK1 is not satisfied for nearly-perfect-homeostasis supporting parameter sets of a four component reaction network with nine reactions and four set point encoding reactions (Fig. 2J). To derive GK2 (Fig. 2H), the weighted dot product between the mass actions of the set-point encoding reactions and sensitivity of the output with respect to the set point encoding rate constants is zero for all perturbations *u* in some range in a perfectly homeostatic system

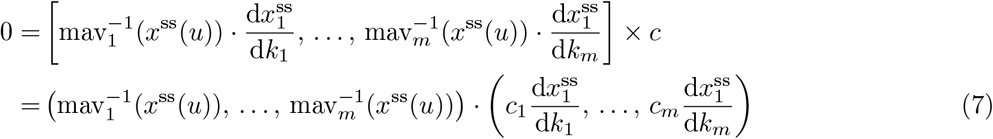

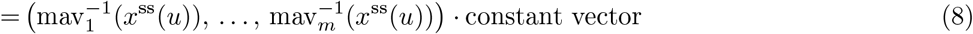

If a non-output system component is also perfectly homeostatic or if two non-output components have the same relative sensitivity, then it is possible for reactant coefficients of non-output components to differ between each two pairs of set-point encoding reactions. This scenario can easily arise if elementary reactions are used to represent transcriptional networks, as many intermediate products leading to the final output of a homeostatic system are all perfectly homeostatic and concentrations of many intermediate products of non-output components are directly proportional to each other. For the four component reaction network with nine reactions and four set point encoding reactions, the proportion of close-to-perfect-homeostasis supporting parameter sets that do not satisfy the generalized GK1 and GK2 conditions is quite high (Fig. 2K).

In summary, existing approaches to find near-homeostasis supporting architectures are not as comprehensive as, or are not faster than brute force search. Some previously reported non-brute force search methods are fast but not comprehensive (*9*), and others are comprehensive but not fast (*2*, 12, 8, 10). Furthermore, these non-brute-force methods only attempt to find perfect homeostasis supporting architectures, which traditionally assumes a simple connection between perfect-homeostasis and near-homeostasis. To translate a perfect-homeostasis supporting parameter set to a near-homeostasis supporting one, the traditional approach is to get the more realistic regulatory dynamics to approach the ideal dynamics of a perfectly-homeostatic system. Saturating proteolytic degradation to approximate zeroth order degradation is one such example (*13, 14*). However, we have previously shown (*2*) and will also show here that near-homeostasis supporting architectures have far more complex properties that cannot be simply translated from perfect-homeostasis supporting architectures. One of many unanswered questions is how much similarity must exist between the realistic regulatory dynamics and the ideal dynamics of a perfectly-homeostasis system for the relative sensitivity to fall below a user-defined threshold (*2*). Thus, near-homeostasis supporting architectures should be discovered and characterized separately.

### Finding near-homeostasis supporting topologies fast using gradient descent within constraints or Lagrangian with rates constraints

We wanted to develop alternative ways to find near-homeostasis supporting architectures that are both comprehensive and faster than brute-force search, and realized that finding near-homeostasis supporting architectures can be recast into a constrained optimization problem. To find a near-homeostasis supporting steady state, we need to identify a point (*p, x*) that lies within the parameter-steady state hyperspace (Fig. 3A: black line) and results in minimal truncated total relative sensitivity as the objective value (Fig. 3A: black *) among all points within that hyperspace. The truncated total relative sensitivity is truncated to the desired threshold *ϵ* > 0 if the total relative sensitivity is below the desired threshold

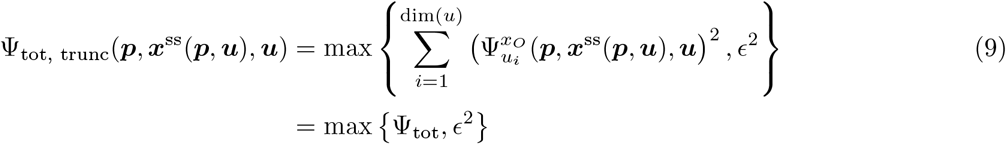

**Fig. 3:**
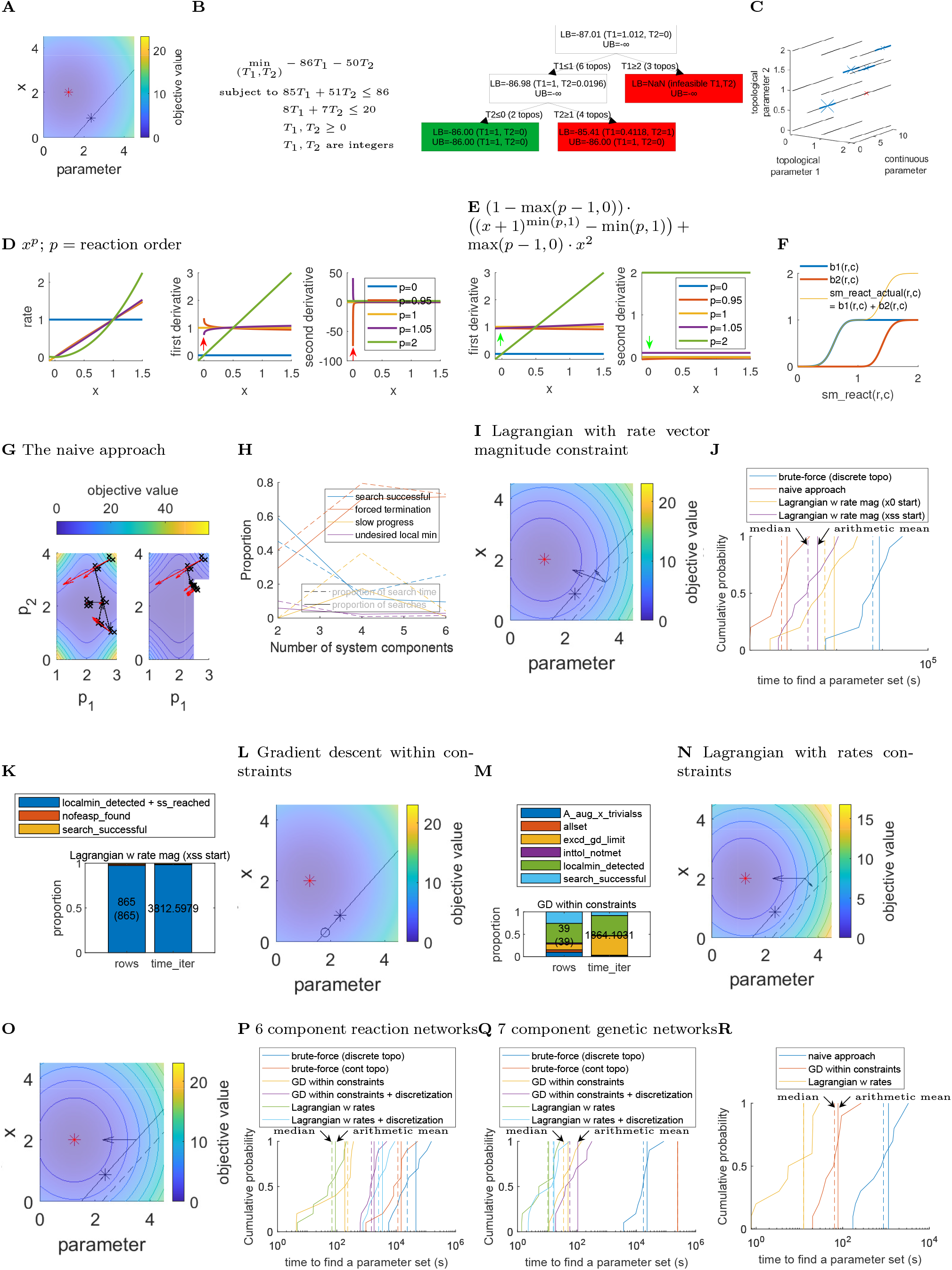
Attempted search strategies to find near-home1o0stasis supporting architectures. ((A) Finding a near-homeostasis supporting parameter set can be cast as a constrained optimization problem. The red* denotes minimum of the objective function. The black line represent the steady state of the system at different parameter values. The black * denotes the parameter steady-state that results in the minimum objective value. (B) Illustration of the branch and bound algorithm to solve mixed integer programming problems. (C) Schematic of finding a near-homeostasis supporting discrete topology. Decreased × size indicates increased consecutive search rounds. The blue line interval denotes near-homeostasis supporting non-topological parameter sets for a fixed topology. The × is red if the search round does not end in a near-homeostasis supporting continuous parameter space. (D) The naive continuous extension of zero, first, and second order reaction rates. Red arrows indicate locations of infinite first and second derivatives. (E) The modified continuous extension of zero, first, and second order reaction rates has continuous first and second derivatives. Green arrows point to the absence of infinite first and second derivatives seen in Fig. 3D. (E) ***b***_**1**_(*r, c*), ***b***_**2**_(*r, c*), and **sm_react_actual**(*r, c*) as functions of the continuous topological parameter **sm_react**(*r, c*). (G) Illustration of the naive approach. × indicate parameters where steady state are simulated. The red arrows indicates empirically determined steepest descent vectors. The black arrows indicate Newton steps taken to find the stationary point of the Lagrange function or equivalently the minimum of the objective function. Unlike the left plot, the right plot contains a parameter region that do not admit steady states. (H) Characterization of the naive approach at increased number of system components. This figure is generated from the classification of searches for near-homeostasis supporting non-topological parameter sets among three increasingly complex reaction networks (Fig. S1). Please see Methods for a detailed explanation of classification labels. (I)Illustration of the Lagrange method with rate magnitude constraints. In addition to Fig. 3A, we also added gradient vectors as black arrows for the objective function and the constraint function. The dashed line is a contour curve for the constraint function. (J) The Lagrange method with rate magnitude constraints is not faster than the naive approach for 2 component reaction networks. (K) Classification of different search attempts for the Lagrange method with rate magnitude constraints. Please see Methods for a detailed explanation of classification labels. (L) Gradient descent within constraints starting from a parameter steady-state coordinate (the circle in addition to Fig. 3A). (M) Classification of different search attempts for the gradient descent within constraints method applied to 6 component reaction networks. Please see Methods for a detailed explanation of classification labels. (N, O) Illustration of the Lagrange method with rates constraints (N) and the Lagrange method with rate vector magnitude constraints (O) starting from a parameter steady-state coordinate. The black arrow(s) in addition to Fig. 3A denote gradient vectors of the objective function and the constraint function at the parameter steady-state coordinate. (P, Q) Comparing the speed of the optimized brute-force method, the gradient descent within constraint method, and the Lagrange method with rates constraints to find near-homeotsasis supporting parameter sets for different types of networks. If a method has the word “discrete” or “discretization”, then that method is asked to find near-homeostasis supporting discrete topologies. (R) The gradient descent within constraints method and the Lagrangian with rates constraints method is faster than the naive approach in finding near-homeostasis supporting continuous topologies of 7 component genetic networks.

The total relative sensitivity is the sum of the squared relative sensitivity of the output at each perturbation coordinate *u*_*i*_. Note that the unconstrained minimum of the objective function (Fig. 3A: red *) does not result in a nearly-homeostatic system, because it fails to yield a steady state for the system. In the rest this section, we characterize the inherent limitations associated with each of the four formulations of the constrained optimization problem, and find two formulations that are able to discover near-homeostasis supporting architectures much faster than brute-force search.

To find a near-homeostasis supporting discrete topology, the most common approaches used in commercial Mixed Integer Programming solvers is the branch-and-bound approach (*15*), and this approach may require extensive sampling of the set of all possible topologies. The branch-and-bound approach (Fig. 3B) considers all possible discretizations of topological parameters and eliminates an entire branch of possible topological values (Fig. 3B: red cell in row 2) whenever the associated subproblem has a minimum objective value that is higher than the objective value of the best performing discrete topology. This approach has exponential worst-case time complexity, where the solution search time can be proportional to 3^*N*×*N*^ when there are *N* system components and every system component can up-regulate, not regulate, or down-regulate any system component (including itself).

However, if we are not concerned with finding the one parameter set with the lowest possible relative sensitivity and instead focus on finding architectures with “good enough” relative sensitivity (as defined by the requirements of a particular control scenario), we can treat the distance of between a continuous topological parameter value supporting near-homeostasis and the nearest discrete topological parameter value as a measure of how likely that discrete value is to also support near-homeostasis. To find a near-homeostasis supporting discrete topology (stage 0), we first start from a continuous topology (Fig. 3C: the largest blue x) and try to find a near-homeostasis supporting continuous parameter set (Fig. 3C: the second largest blue x). In the next stage (stage 1), we discretize topological parameters that are sufficiently close to a discrete state (Fig. 3C: the third largest blue x). For topological parameters not close enough to a discrete state, we randomly assign them to a possible discrete value and favor the closest discrete value in stage 2. Multiple short stage 2 discretization iterations can be carried out to find a near-homeostasis supporting discrete topology (Fig. 3C: the smallest blue x), as not all adjacent discrete topological states enable near-homeostasis (Fig. 3C: red x).

The continuous topological parameter extension should have continuous transitions between different topological values for the zeroth, first, and second derivatives; it must also be identical to the original network having a discrete topology when the topological parameters are discrete. For reaction networks described in Fig. 2F, if the general form of the mass action is *x*^*p*^ as one might expect and the topological parameter is close to 1, then the first and second derivative of the continuous extension diverges to infinity for *x* near zero (Fig. 3D). Thus, we have developed a more complicated continuous extension of the mass action consumption of *x* that guarantees a continuous transition between different topological states for the zeroth, first, and second derivative (Fig. 3E). However, the derivative of min(*p* − 1, 0) with respect to *p* is discontinuous rather than having the continuity property we seek, and we also want to slow down changes in topological parameters relative to non-topological parameters whenever the former are nearly-discrete. Therefore, we further modify the mass action of *x*_*r*_ in reaction *c* (mas(*x*_*r*_, *r, c, b*_3_)) to satisfy the additional two criteria

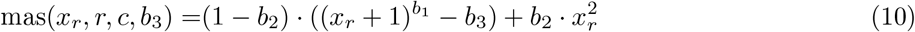

Here, *b*_1_ and *b*_2_ are smoothed step-functions of the continuous topological parameter **sm_react**(*r, c*) (Fig. 3F) that attain fully discrete mapped topological values

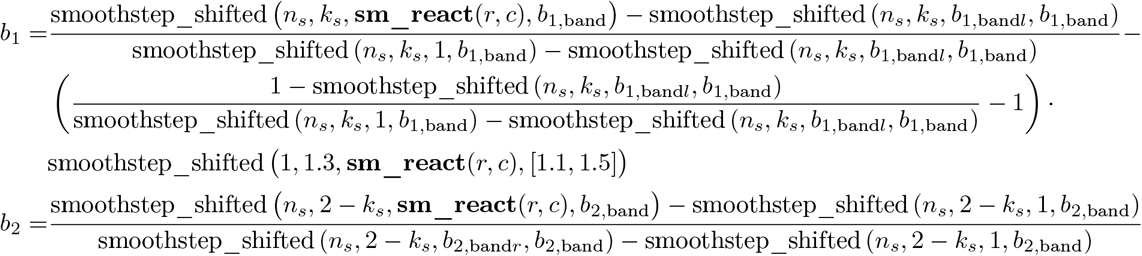

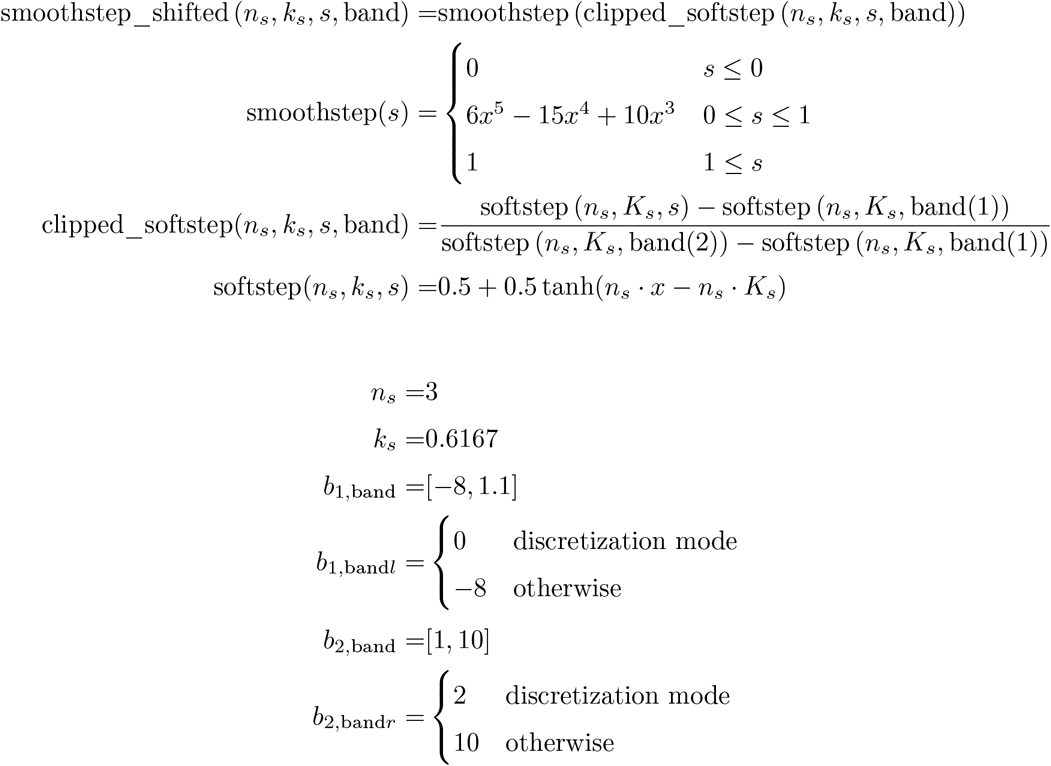

The continuous extension of a two component reaction network is then

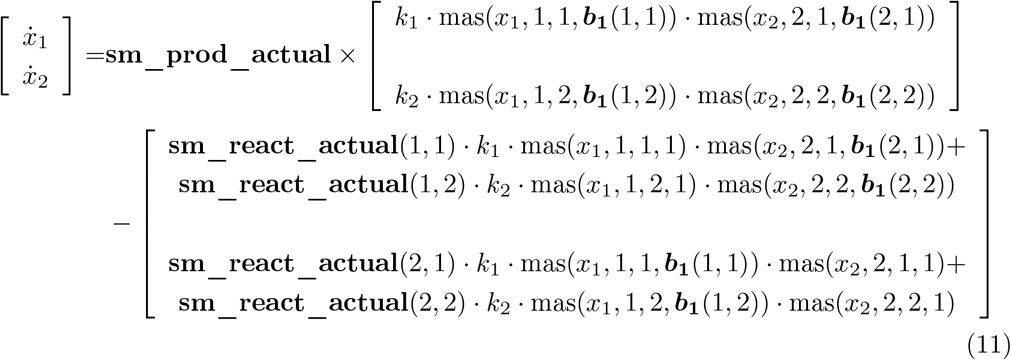

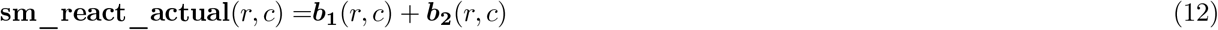

where the flux vector changes when multiplied with different rows of the reactant stoichiometric matrix sm_react. If **sm_react_actual**(1, 1) = 0.5 and the flux vector does not change when multiplied with different rows of the reactant stoichiometric matrix in a two component elementary reaction, then the flux multiplying **sm_react_actual**(1, 1) in the first row surprisingly does not result in 0 at *x*_1_ = 0, *x*_2_ > 0

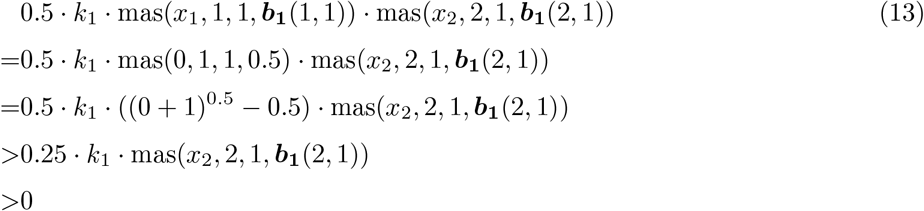

When all stoichiometric coefficients are non-negative integers, the continuous extension Eqn 11 returns identical rates as the discrete mass of action network in Fig. 2F.

One naive formulation is to simulate the steady state and compute the truncated total relative sensitivity at multiple continuous topological and non-topological parameters (Fig. 3G: left plot, crosses) to figure out the direction of parameter change that minimizes the truncated total relative sensitivity to the greatest extent (Fig. 3G: left plot, red arrows, also called gradient vectors) and use root finding algorithms to find parameter sets that result in zero gradient vectors (Fig. 3G: left plot, black trajectory), but this approach becomes increasingly cumbersome and failure prone as network size increases. An approach along the same line of reasoning was suggested by the large language model GPT-4 (*16*). One major drawback of this approach is that the number of steady state simulations required to determine each gradient vector increases with the number of parameters, and the number of parameters increases rapidly (usually quadratically) as the number of network components increases. The second major drawback of this approach is that the search algorithm often converges to a region of parameter space where no parameter set admits a steady state (Fig. 3G: right plot, white region), resulting in forced termination of the parameter search. As network size increases, the proportion of forced termination events quickly increases and the proportion of successful search results quickly decreases (Fig. 3H). Here, this approach has already attempted to minimize forced terminations by incorporating the ability to backtrack away from an encountered no-steady-state parameter region.

Another technique is to use root finding algorithms to find stationary points of the Lagrangian function with the magnitude of the rate vector as the constraint

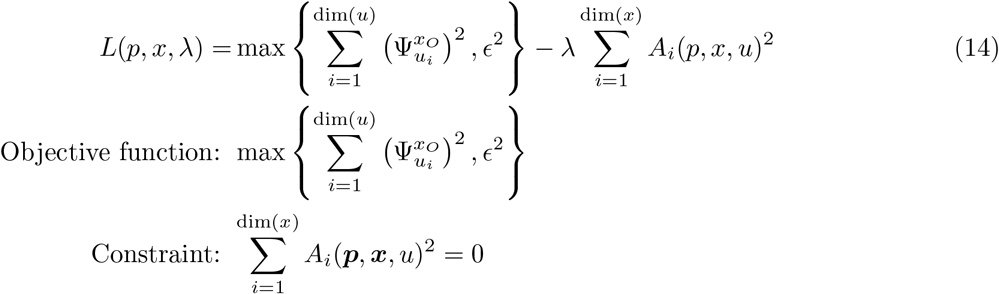

A stationary point of the Lagrangian function does not lie on a slope in any given direction

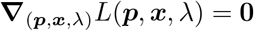

At non-stationary points of the Lagrange function, the object function’s gradient vector and the constraint function’s gradient vector are not parallel or the constraints are not satisfied. If the two gradient vectors are not parallel, the contour of the objective function would cross the contour of the constraint function and there exists a direction to walk within the contour of the constraint function such that the objective function value decreases (Fig. 3I). Thus, the desired near-homeostasis supporting parameter sets result in the objective function’s gradient vector and constraint function’s gradient vector being parallel to each other and the nonlinear constraint being satisfied, which are precisely the stationary points of the Lagrange function. Unfortunately, this algorithm is slower than the naive approach for multiple systems (Fig. 3J: two component reaction network; Fig. S2: Ma et al’s negative feedback loop). The biggest issue with this approach is that even for simple networks, most of the search time is spent in a state of near-constant total relative sensitivity but the magnitude of the rate vector is near zero (Fig. 3K; Fig. S2).

We tried a third approach to find near-homeostasis supporting architectures by walking within the parameter-and-steady-state hyperplane down the steepest parameter directions that decrease the total relative sensitivity (Fig. 3L), which we call gradient descent within constraints. We created an augmented dynamical system that combines the native dynamics of the original system components (cyan highlights), how much the steady state of the system changes in response to a certain amount of change in parameter values (red highlights), and how much parameter values should change to result in the steepest decrease in the total relative sensitivity while the system components are always at steady state (yellow highlights).

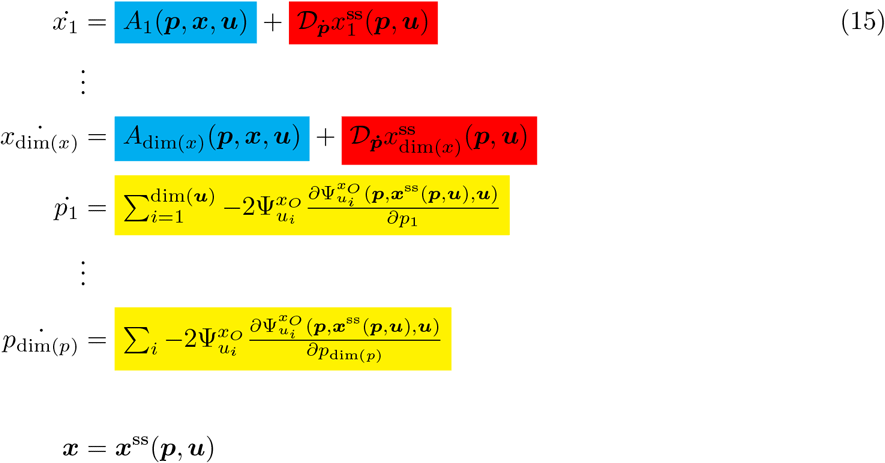

The coupling between the system state variables and the parameter values ensures that the augmented system’s trajectories remain within the parameter-and-steady-state hyperplane rather than potentially drifting either into regions of parameter space where no system steady state exists, or into system steady states far from the optimal regions of parameter space. The parameter changes required to result in the steepest decrease in total relative sensitivity are different depending on whether the system components must always be at steady state. In the example illustrated in Fig. 3L, consider starting from a point on the parameter-and-steady-state hyperplane (Fig. 3L: black circle): from that point, increasing the parameter decreases the objective function value towards the black star given the constraint of remaining within the parameter-and-steady-state hyperplane. On the other hand, if no hyperplane constraint were present, the objective function could be reduced further by decreasing the parameter value from the black circle to the red star. An important component of this algorithm is using automatic differentiation to minimize the number of reevaluations. For instance, the most expensive operation in computing parameter gradients is computing the second derivative of the Jacobian matrix with respect to each parameter:

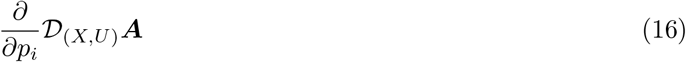

For a two component reaction network with one reaction (Eqn 11), computing the second derivative starting from fully substituted expressions involves repeated computation of the same expressions (highlighted in cyan and red):

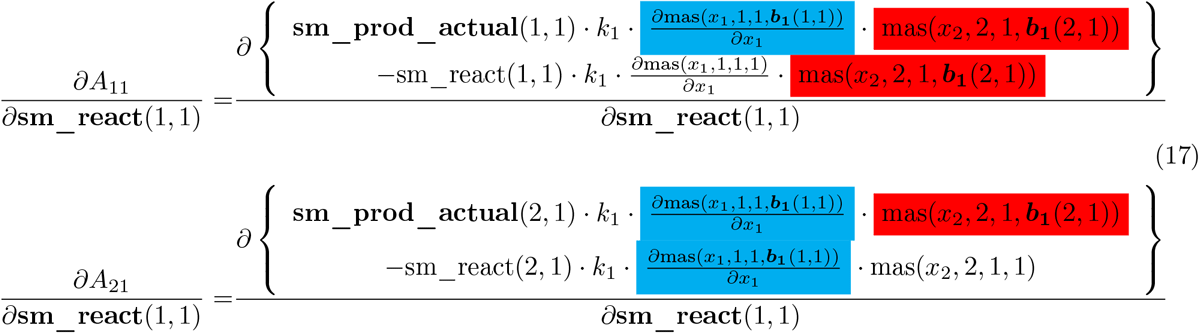

Automatic differentiation not only enables computation of arbitrary derivatives, it also ensures that the derivative of a repeatedly used expression is only computed once. Such repeatedly used expressions are much more frequently occurring in our case than during back-propagation of neural networks, making automatic differentiation even more crucial. For complex networks, the progress of each search can be somewhat slow and more than half of the search time is wasted on searches where the total relative sensitivity is trapped above the desired threshold (Fig. 3M: localmin_detected). However, for these networks, this approach allowed us to find near-homeostasis supporting topologies at least 10 times faster than brute search (Fig. 3P, 3Q).

In an attempt to address the sometimes slow search progress of the gradient descent within constraints approach, we tried to find a stationary point of the Lagrangian that minimizes total relative sensitivity but subject to every rate equation being zero rather than the magnitude of the rate vector being zero

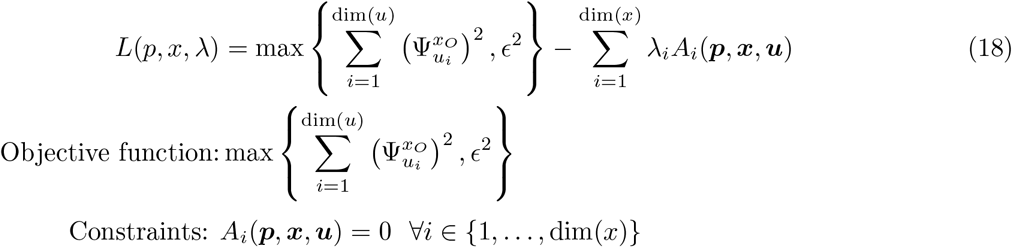

This variant seems quite similar to the other Lagrange optimization approach, but non-parallel gradient vectors occur in slightly different scenarios, highlighting the importance of understanding the intuition behind Lagrange multipliers. However, when the set of all rate equations are used as the nonlinear constraints, the Lagrange algorithm would be able to detect “non-parallel” gradients vectors of the relative sensitivity function and of the set of all rate equations for points on the steady state curve (Fig. 3N). At points above relative sensitivity threshold but on the steady state curve itself, the gradient vector of the magnitude of rate vector is zero and is automatically parallel to the gradient vector of the objective function (Fig. 3O). Similar to the gradient descent within constraints approach, automatic differentiation was also implemented to vastly minimize the number of reevaluations. Although this algorithm is much faster than brute-force search at finding near-homeostasis supporting discrete topologies (Fig. 3P, 3Q), the algorithm is not significantly faster than the gradient descent within constraints approach. Lastly, both the gradient descent within constraints approach and the Lagrangian with rates constraints approach are faster than the naive approach for sufficiently complex systems, often orders of magnitude faster (Fig. 3R, S3).

It is important for our two successful optimization algorithms to be fast at finding near-homeostasis supporting topologies for multiple types of regulatory networks: elementary reaction networks and higher level transcriptional networks. While elementary reaction networks can theoretically represent any biological regulatory network, higher level descriptions like Hill functions often vastly reduce the number of rate equations needed to represent the network compared to using elementary chemical reactions (*17*). Unlike reaction networks, each system component in a transcriptional network can up-regulate, down-regulate, or not regulate any system component and its regulatory effect saturates at a sufficiently high concentration

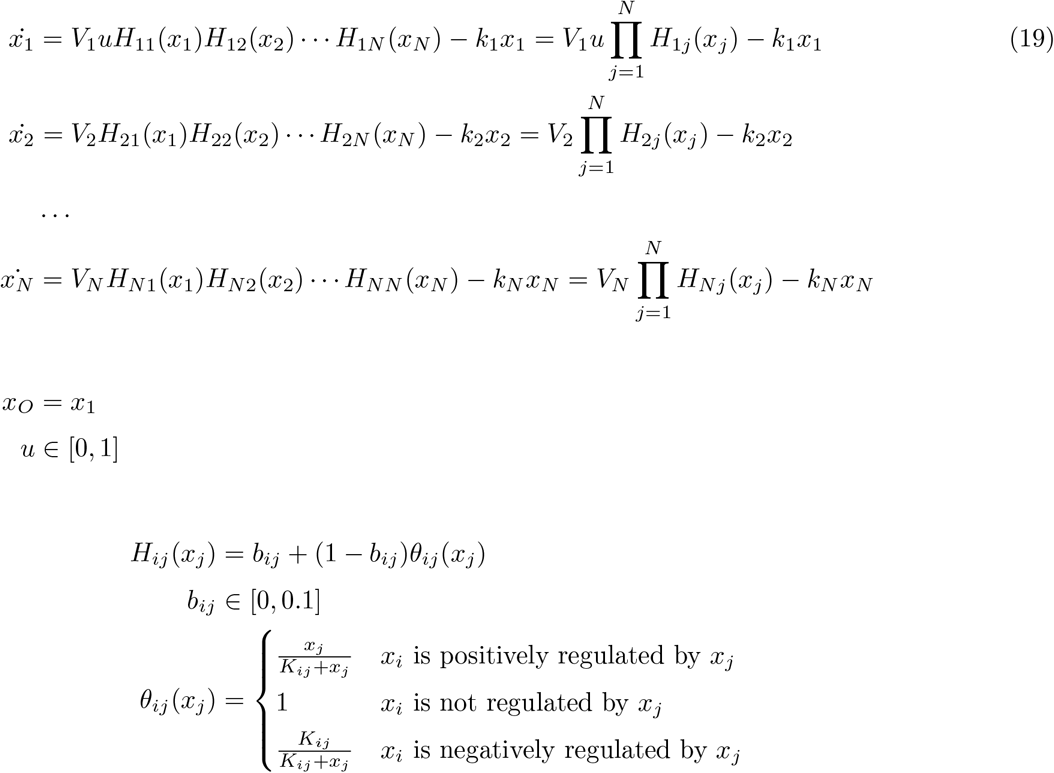

Here, *x*_1_ is the system output and *u* is the scalar perturbation. As topological parameter for the regulation of *x*_*i*_ by *x*_*j*_ (*s*_*ij*_) increases, the continuous extension of *θ*_*ij*_

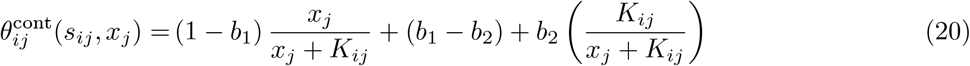

switches continuously from positive regulation at *s*_*ij*_ = 0 to no regulation at *s*_*ij*_ = 1 and finally to negative regulation at *s*_*ij*_ = 2. The definition of *b*_1_, *b*_2_ are identical to the chemical reaction network’s general form (Eqn 10).

### Traditional characterization of near-homeostasis mechanisms is incomplete and requires complicated statistical analysis

Once enough near-homeostasis supporting parameter sets are found, we must then characterize commonly occurring mechanisms of near-homeostasis. Each parameter set supporting near-homeostasis comes as a list of numerical values. Because precise parameter values cannot be implemented in biological systems due to uncertainties surrounding their implemented values (*18, 19*), we must sufficiently understand the mechanism responsible for near-homeostasis to perform the experimental implementation. Furthermore, understanding the mechanisms underlying near-homeostasis would allow us to compare their respective advantages given the experimental constraints. In this section, we demonstrate that traditional characterization of nearhomeostasis mechanisms (*1,5*) requires a great deal of intuition and complicated statistical analysis, and still cannot comprehensively identify near-homeostasis mechanisms.

The difficulties encountered by the traditional approach become apparent when we want to discover the mechanisms of near-homeostasis for a collection of incoherent feedforward loops, where *x*_3_ is the output (Fig. 4A). Between near-homeostasis supporting parameter sets and those that result in large sensitivity to perturbation, we can compare the distributions of any of the many parameters or any of the system component steady states like *K*_21_ and *x*_2_ (Fig. 4B). We can then propose composite metrics that are predicted and may later be validated to be differentially distributed between instances of successful near-homeostatic performance and instances of failure to maintain near-homeostasis. Designing suitable composite metrics is not very intuitive and relies on a high amount mathematical intuition. In this case, the maximum of the quantile for *K*_21_*/*(1 − *x*_2_) and quantile for 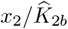 is somewhat differentially distributed between nearly-homeostatic and not-homeostatic parameter sets. Indeed, if we assume those two ratios are sufficiently small, then the *x*_2_ rate equation enforces an approximately linear relationship between *x*_1_ and *x*_2_ at steady state (Fig. 4C). That approximately linear relationship allows the third rate equation to specify a nearly perturbation independent *x*_3_ steady state, by making *x*_2_ steady state separable. However, this previously described mechanism of near-homeostasis is only possessed by a fraction of near-homeostasis supporting parameter sets. For the majority of near-homeostasis supporting parameter sets, the maximum of the quantile for *K*_21_*/*(1−*x*_2_) and quantile for 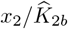 are above the 5th or even 10th percentile of not-homeostatic parameter sets.

**Fig. 4:**
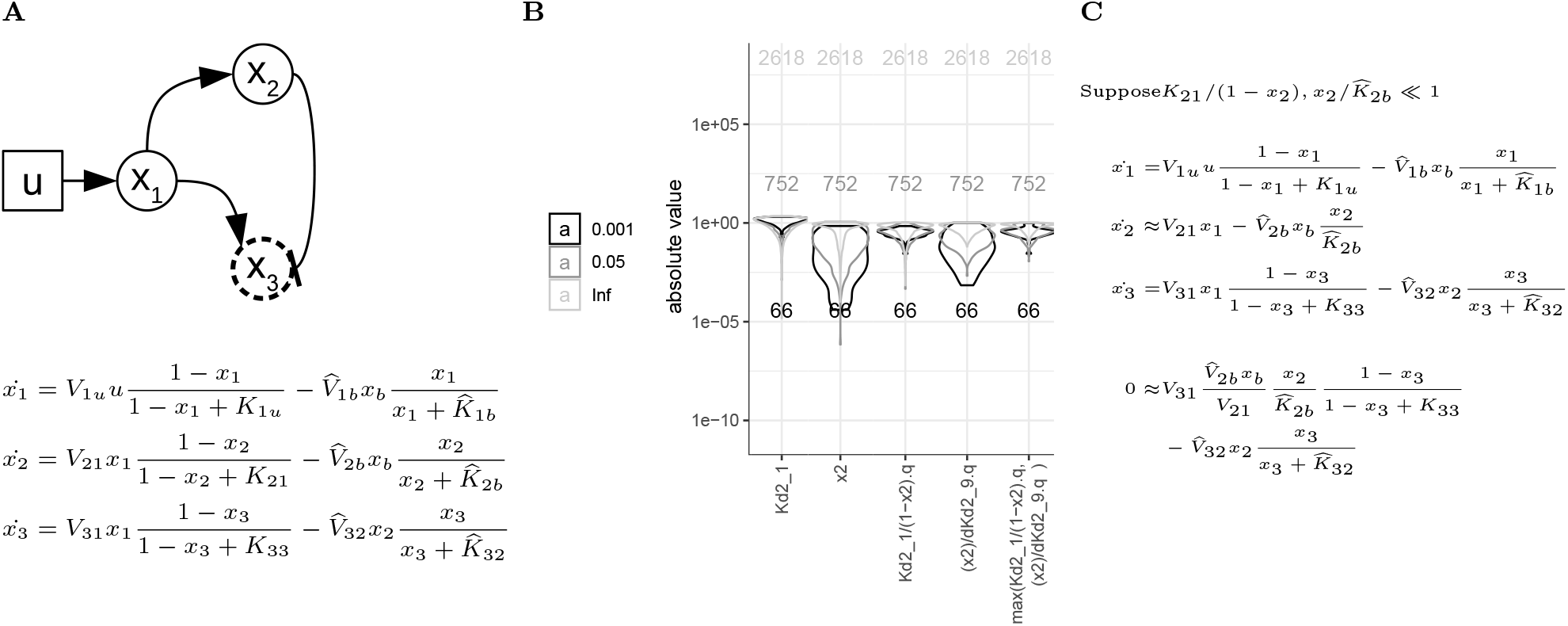
Trial-and-error based statistical analysis of parameter sets associated with incoherent feedforward loops. (A) Schematic and mathematical description of incoherent feedforward loops. (B) Comparing near-homeostasis supporting parameter sets that has maximal relative sensitivity of 0.001 against not far-from-homeostatic parameter sets that has minimal relative sensitivity of > 0.05, in terms of different user-defined metrics. (C) Ma et al (*1*)’s mechanism of near-homeostasis for the incoherent feedfoward loop.

### Inverse homeostasis plots are used to identify novel mechanisms of near-homeostasis without using complicated statistical analysis

We wanted to identify common mechanisms of near-homeostasis for five component transcriptional networks (Eqn 19: N=5), and discovered a graphical approach to easily and comprehensively characterize those common mechanisms. We realized that all mechanisms of near-homeostasis can be understood by restating the problem in terms of how a small fold difference in steady state output is induced by large changes in steady states of other key components and how such large steady state changes is ultimately induced by a large change in perturbation (Fig. 5A). We call this approach the inverse homeostasis perspective, which inverts the usual approach of analyzing how a large change in perturbation results in a small fold change in output steady state. Using our approach, we discovered a new two-component nearly-homeostatic controller that tolerates much higher undesired basal expression levels compared to the well-known one-component autocatalytic controller.

**Fig. 5:**
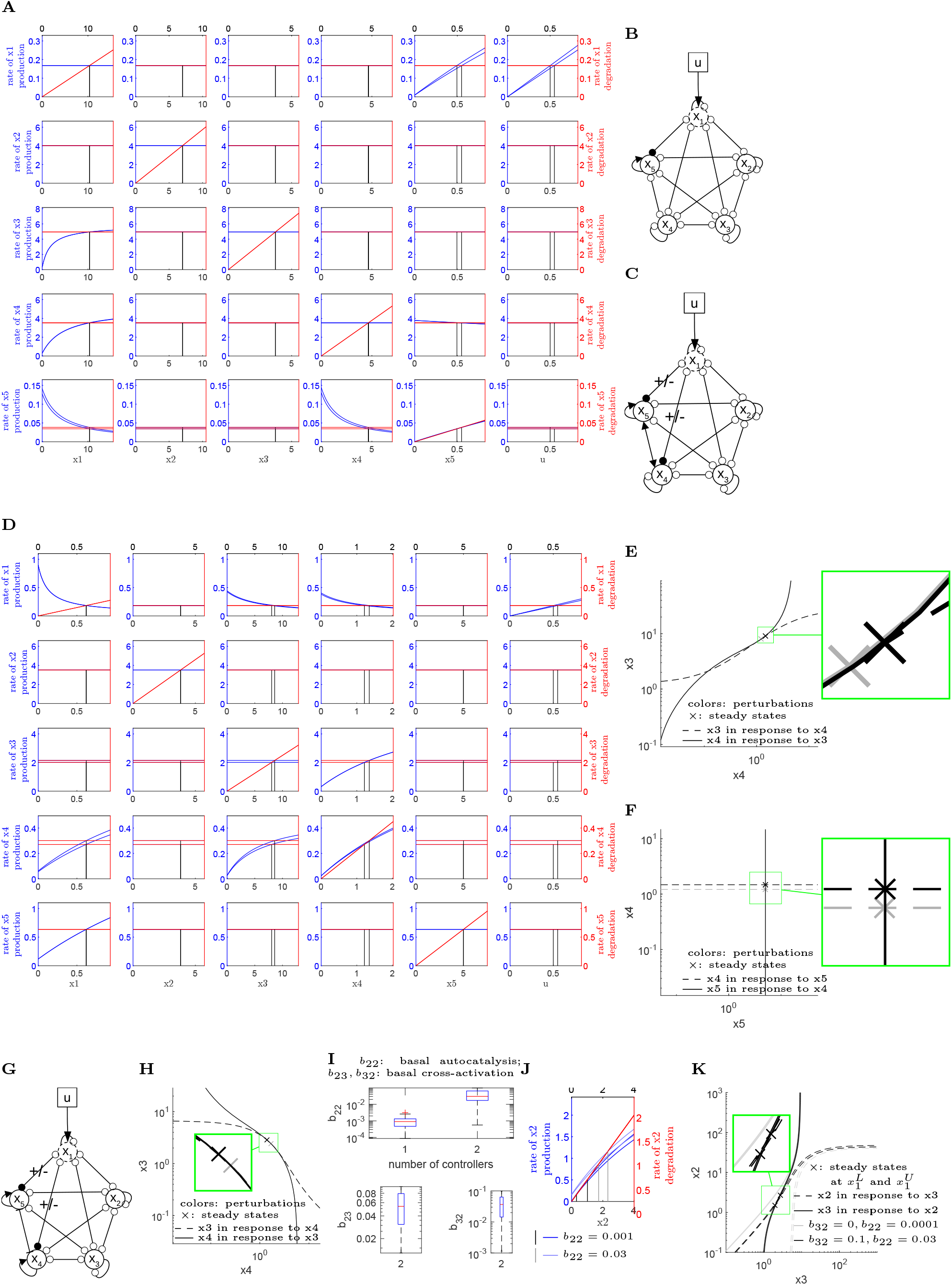
Characterization of near-homeostasis supporting parameter sets for 5 component transcriptional networks. (A) Level 1 inverse homeostasis plot of an autocatalytic nearly-homeostatic system. Vertical black lines are the anchoring steady states that were used to generated the level 1 inverse homeostasis plot. (B) Topology of autocatalytic nearly-homeostatic systems. (C) Topology of cross-activation based nearly-homeostatic systems. (D, E, F) Level 1 (D) and level 2 (E, F) inverse homeostasis plot of a cross-activation based nearly-homeostatic system. Here, *x*_3_ and *x*_4_ activate each other. For level 2 inverse homeostasis plots, the steady state curves of *x*_*i*_ in response to *x*_*j*_ at different perturbations are often nearly identical to each other, because the perturbation affects neither system components {*x*_*i*_, *x*_*j*_} and only minimally changes the system output steady state. (G)Topology of cross-repression based nearly-homeostatic systems. (H) Level 2 inverse homeostasis plot of a cross-repression based nearly-homeostatic system. Here, *x*_3_ and *x*_3_ repress each other. (I) Comparing near-homeostasis supporting parameters of the single component autocatalytic controller against the two component cross-activation controller. (J) Level 1 inverse homeostasis plot for the single component autocatalytic controller at different basal autocatalysis rates. (K) Level 2 inverse homeostasis plot for the two component cross-activation controller at different basal expression rates.

To generate the first level of plots for the inverse homeostasis perspective, we plot the production rate and degradation rate of every system component against every system component and every perturbation, using the steady states of a nearly-homeostatic system at all perturbations as anchor points. For example, to plot the rate of production of *x*_1_ against *x*_5_, we take the steady state of the system at the lower end of the perturbation range *x*^ss^(*u*^*L*^), change the *x*_5_ component, and plot the corresponding *x*_1_ production rate as the lower blue line (Fig. 5A). We then do the same for the steady state of the system at the upper end of the perturbation range *x*^ss^(*u*^*U*^) to plot the upper blue line. We then plot the degradation rate of *x*_1_ against *x*_5_ as red lines in Fig. 5A, using steady states of the system at lower and upper perturbations. In this case, since the degradation of *x*_1_ does not depend on *x*_5_, both degradation rate curves are horizontal and identical. Each anchor steady state results in intersection of the production rate curve and degradation rate curve.

We then find a sequence of level 1 inverse homeostasis plots that explain how a very small fold change in steady state output *x*_1_ is induced by a relatively large fold change in perturbation *u*. Although there are dim(***x***) × (dim(***x***) + dim(*u*)) inverse homeostasis plots corresponding to the permutations of every state with all states and all inputs, we can immediately ignore plots where the steady states are as close together as the system output or both production and degradation rate curves are horizontal. For the five state, one input system under consideration, there are 5 × 6 grid of level 1 inverse homeostasis plots (Fig. 5A). Based on the aforementioned criteria, we can dismiss from detailed consideration plots (2:4,1:4), (2:3,5:6), (4,6), (5,2:3), and (5,6) [(row,column), with the range from *i* to *j* indicated by (*i*:*j*)]. According to plot (5,5), a very small fold increase in the output *x*_1_ steady state and *x*_4_ steady state would result in a slight decrease in *x*_5_’s production rate curve. The slight decrease or downward rotation of *x*_5_’s production rate curve is sufficient to induce a relatively large fold decrease in *x*_5_ steady state. The very small fold change in the output *x*_1_’s steady state results in a very small fold change degradation rate of *x*_1_ at steady state (Fig. 5A: plot (1,1)). The relatively large fold decrease in *x*_5_ steady state combined with the very small fold change in the output *x*_1_’s steady state results in two diverging production rate curves (Fig. 5A: plot (1,6)). The two diverging production rate curves intersect with the two nearly identical steady state degradation rates to result in a relatively large fold increase in perturbation *u*. The only topological requirement is that a nearly-homeostatic component must directly regulate the autocatalytic controller (Fig. 5B). This single-component autocatalytic mechanism of near-homeostasis has been reported in different previously analyzed networks (*1*, 20, 9).

We have further discovered some new mechanisms of near-homeostasis that are better explained by intersections of the steady state curves of two system components in response to one another, called level 2 inverse homeostasis plots. For the near-homeostasis supporting parameter set we will consider here, there are two controller components that cross-activate each other, at least one of two controllers has autocatalysis, and a nearly homeostatic component directly regulates one of the two controllers (Fig. 5C). When the nearly-homeostatic component directly regulates both controllers, the type of regulation is the same in almost all cases. According to the level 1 inverse homeostasis plots generated for this parameter set, a very small fold change in the output *x*_1_ at steady state results in a relatively large fold change in *x*_3_ and *x*_4_ steady state, but it is difficult to tell how the large fold change in those two system components came about (Fig. 5D). However, it is very interesting that the subsystem created by *x*_3_ and *x*_4_ permits two very different steady states at two nearly-identical output *x*_1_ steady states and is independent of perturbation *u* (Fig. 5D: plots (3:4,3:4)). Thus, for each pair (*x*_*i*_, *x*_*j*_), *i* ≠ *j* of the non-system output components {*x*_2_, …, *x*_5_}, we plot two steady state curves of *x*_*i*_ in response to *x*_*j*_, using as anchor points the steady state in the presence of the lower perturbation (Fig. 5E: gray curve) and the steady state in the presence of the upper perturbation (Fig. 5E: black curve). At each anchor steady state, changing *x*_*j*_ almost always results in a unique *x*_*i*_ steady state. We also plot two steady state curves of *x*_*j*_ in response to *x*_*i*_ using the same two anchoring steady states. This phase diagram of *x*_*i*_ and *x*_*j*_ subsystem has four curves, two corresponding to the response of *x*_*i*_ toward *x*_*j*_ at the two fixed perturbations and another two corresponding to the response of *x*_*j*_ to *x*_*i*_ at the two fixed perturbations. Here, a small fold change in *x*_1_, *x*_2_, *x*_5_ at steady state results in a very slight shift in the response of *x*_4_ toward *x*_3_ (Fig. 5E: nearly-overlapping solid curves), which is sufficient result in a relative large fold change in *x*_3_ and *x*_4_ steady state. In contrast, the steady response curves between *x*_4_ and *x*_5_ is uninformative, as it only shows how a small fold change in *x*_1_, *x*_2_ but a relatively large fold change in *x*_3_ results in a relatively large fold change in *x*_4_ steady state (Fig. 5F). This mechanism of near-homeostasis will be called two-component cross-activation with autocatalysis.

For five component genetic networks, most near-homeostasis supporting parameter sets can be explained using level 1 or level 2 inverse homeostasis plots. Only 4 out of 25 near-homeostasis supporting parameter sets could not be explained using either level 1 or level 2 inverse homeostasis plots (Supplementary Information GN_unknown_uniqdeg-nhdtp_focus-characterization.xlsx). The three common mechanisms of near-homeostasis are the single-component autocatalysis (Fig. 5A, 5B), the two-component cross-activation with autocatalysis (Fig. 5C, 5E), and two-component cross-repression with autocatalysis (Fig. 5G, 5H).

Importantly, two component cross-activation with autocatalysis tolerates much higher basal expression than a single component autocatalytic controller (Fig. 5I). To ensure a fair comparison between the two controller types, only one of the two components is autocatalytic for the two component cross-activation with the autocatalysis controller

One component autocatalytic controller

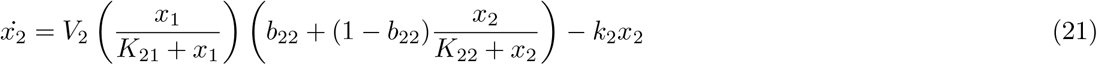

Two component cross-activation controller with autocatalysis

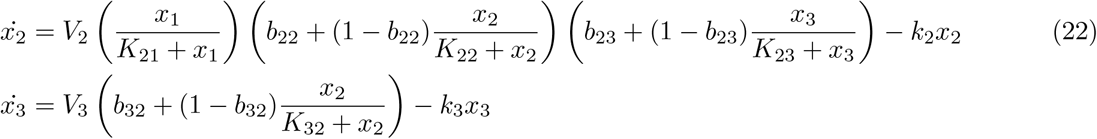

For each of the cross-activation with autocatalysis controller and the one component autocatalytic controller, we randomly found 50 parameter sets such that all controller components had ≥ 6 fold higher log unit change in steady state for a fixed change in output. According to the inverse homeostasis plots, a fixed fold change in output resulting in a much larger fold change in controller steady state is necessary for near-homeostasis. Such simpler criterion can therefore be used to compare the potential of different controllers for near-homeostasis, independent of non-controller components within the feedback system that are difficult to standardize between single-component and dual component controllers. We then compared the distribution of basal expression rates, and found 10^1.5^ = 31.6 fold higher tolerance for basal autocatalytic expression *b*_22_ for the cross-activation + autocatalysis controller with little restriction on basal cross-activation *b*_23_, *b*_32_ beyond its original range of [0, 0.1]. For the single component autocatalytic controller, very small basal autocatalysis *b*_22_ enables a good approximate overlap between linear region of the controller’s production rate curve and degradation rate curve at a suitable output value, allowing a small fold change in output to result in a large fold change in controller steady state (Fig. 5J: blue curves and black vertical lines). High basal autocatalysis delays the range of controller values for which good approximate overlap is possible, so there is a large reduction in the maximal fold change in controller steady state in response to a small output change (Fig. 5J: light blue curves and gray vertical lines). For the cross-activation controller on the other hand, higher basal cross-activation *b*_32_ shifts the steady state curve of *x*_3_ in response to *x*_2_ to higher *x*_3_ levels and makes the curve less responsive to *x*_2_ (Fig. 5K: solid black curve vs solid gray curve), but sufficiently high basal autocatalysis *b*_22_ lowers sensitivity of *x*_2_ steady state in response to *x*_3_ such that a small fold change in *x*_1_ results in a large fold change in *x*_2_ and *x*_3_ steady state (Fig. 5K: dashed black curve vs dashed gray curve).

### Inverse homeostasis plots are used to rationally modify the antithetic integral controller to truly attenuate the detrimental effect of dilution

We also wanted to see whether our search algorithm and inverse homeostasis plots would allow us to discover new classes of near-homeostasis supporting architectures for multi-perturbation reaction networks that had first been introduced by Gupta et al. (*9*). Successful identification of near-homeostasis supporting architectures for reaction networks demonstrates versatility of our approach. We explored multi-perturbation reaction networks in ways that are well outside the scope of Gupta et al. (*9*)’s search algorithm. First, we allowed perfectly homeostatic set-point values to be encoded by three rather than two elementary reactions. Second, we looked for near-homeostasis supporting parameter sets in addition to those supporting perfecthomeostasis. We identified some of the same near-homeostasis mechanisms found in nearly-homeostatic genetic regulatory networks. Furthermore, level-two inverse-homeostasis plots of the well known antithetic controller inspired a novel modification that made the new antithetic controller truly resistant to the detrimental effect of first-order dilution.

Similar to genetic regulatory networks, near-homeostasis supporting parameter sets for reaction networks can have autocatalysis (Fig. 6A: row 3), cross-activation (Fig. 6B), and cross-repression near-homeostasis mechanisms (Fig. 6C). In particular, we note that the antithetic integral controller (*6, 21*) is an instance of the cross-repression controller. At steady state, the two controller components of the antithetic integral controller

**Fig. 6:**
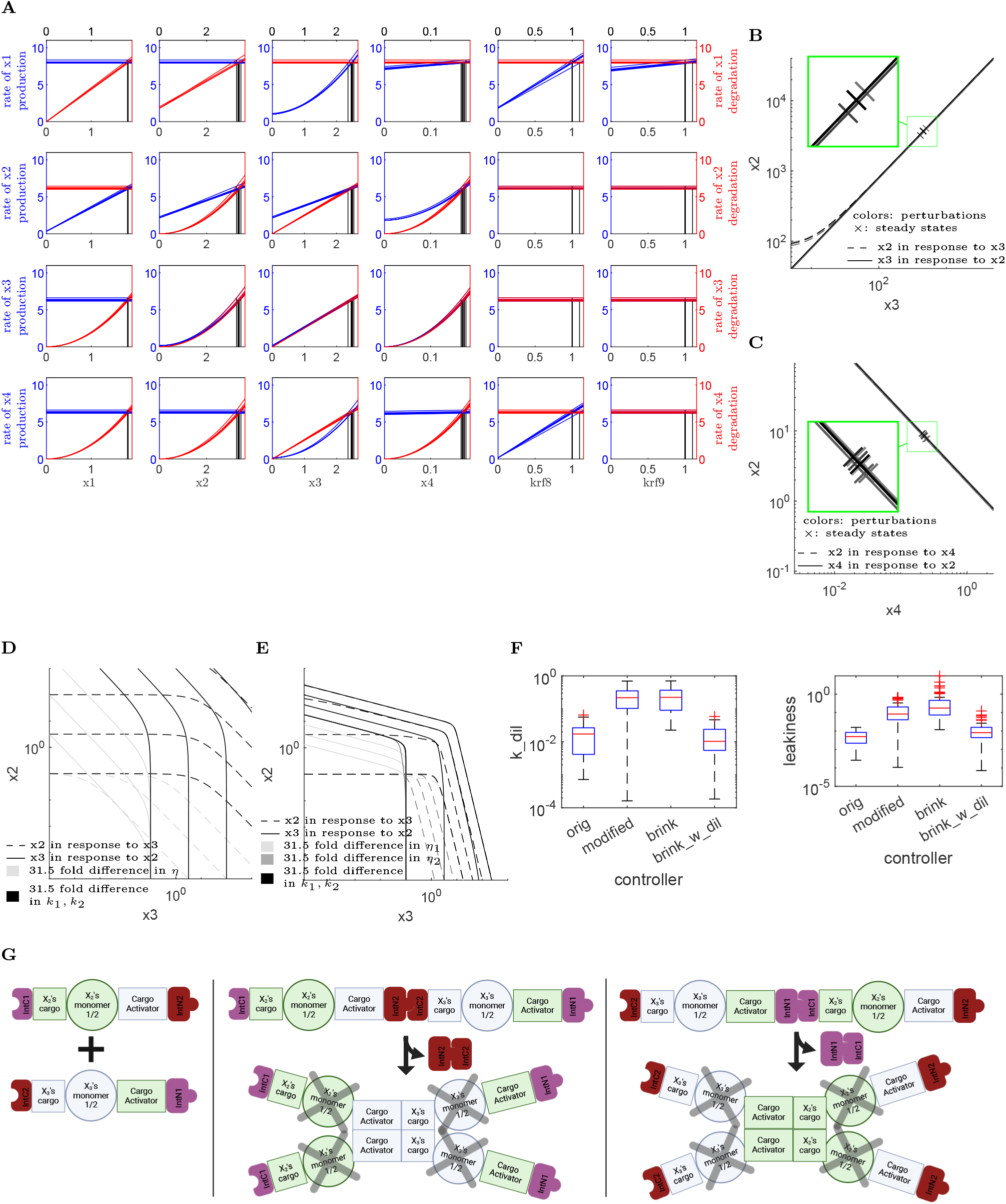
Improving the antithetic controller by characterizing near-homeostasis supporting parameter sets of 4 component reaction networks. (A) Level 1 inverse homeostasis plot of a autocatalytic nearly-homeostatic reaction network. (B) Level 2 inverse homeostasis plot of a cross-activating nearly-homeostatic reaction network. (C) Level 2 inverse homeostasis plot of a cross-repressing nearly-homeostatic reaction network that the antithetic controller (*6, 21*) is a member of. (D) Level 2 inverse homeostasis plot of an antithetic controller at different parameters. (E) Level 2 inverse homeostasis plot of the modified antithetic controller at different parameters. (F) Comparing near-homeostasis supporting parameter sets of the antithetic controller, the modified antithetic controller, the brink controller, and the brink controller with a dilution term for all species. (G) Implementation schematic of the modified antithetic controller using intein-gated dimerization-based inactivation of the controllers’ active domains.

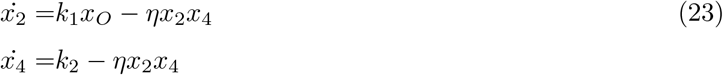

has identical steady state curve in response to each other and that curve has a slope of −1 on the log scale

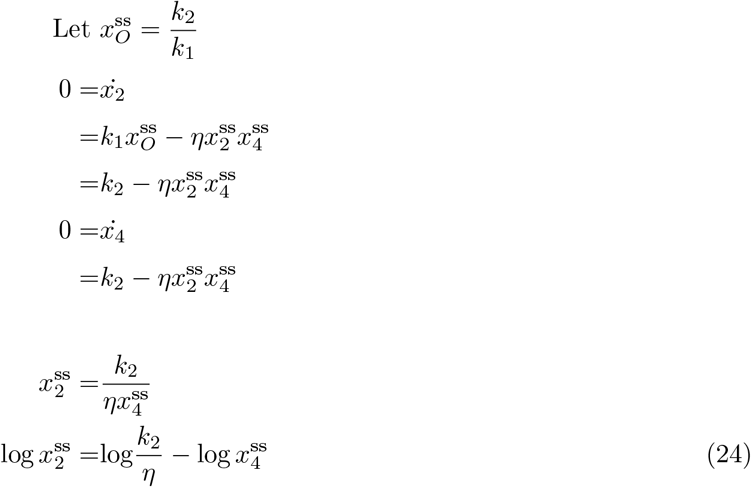

The parameter *η* simultaneously changes both the steady state curve of *x*_2_in response to *x*_4_ and the steady state curve of *x*_4_ in response to *x*_2_, such that the two curves still maintain near-complete overlap with each other.

While prior characterizations of the antithetic integral controller have studied the effect of dilution using algebraic approaches (*22, 7*), our inverse homeostasis plots show why dilution *k*_dil_ within the antithetic integral controller

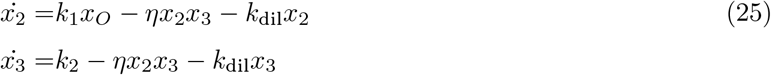

is quite detrimental to near-homeostasis in a far more intuitive manner. In the presence of dilution, the steady state curve of *x*_2_ in response to *x*_3_ has a slope between (−1, 0) on the log scale and the steady state curve of *x*_3_ in response to *x*_2_ has a slope between (−∞, −1) on the log scale

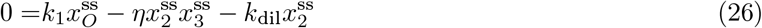

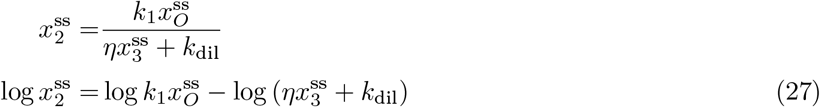

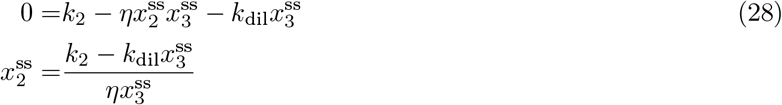

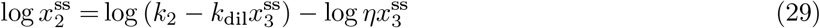

In particular, the slope ranges of the steady state response curves do not overlap (Fig. 6D), but increasing the controllers’ production rate constants *k*_1_, *k*_2_ (Fig. 6D: black curves) and increasing the annihilation rate constant *η* relative to the dilution rate constant *k*_dil_(Fig. 6D: gray curves) by 10^1.5^ ≈ 31.6 fold at a time each enlarged the region of similar slopes between the two response curves. We define integration leakiness, a notion first introduced by Qian et al. (*23*), as relative amount of dilution in relation to the production rates and annihilation rates

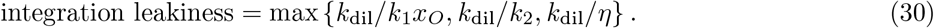

In other words, integration leakiness needs to be quite low in order to achieve near-homeostasis.

We hypothesized that making the slope ranges of the two controllers’ steady state response curves overlap might increase the integration leakiness tolerance that maintains near-homeostasis, by keeping the same region of similar slopes between the two response curves at higher integration leakiness. The modified antithetic controller we propose,

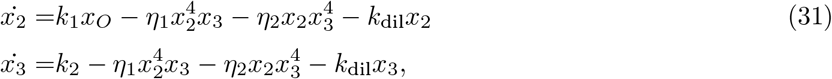

has a slope range of (−4, 0) for the steady state curve of *x*_2_ in response to *x*_3_ and an overlapping slope range of (−∞, −1*/*4) for the the steady state curve of *x*_3_ in response to *x*_2_ (Fig. 6E). Increasing production rates *k*_1_, *k*_2_ (Fig. 6E: black curves) and increasing annihilation rates *η*_1_, *η*_2_ (Fig. 6E: light gray and dark gray curves) by 10^1.5^ ≈ 31.6 fold at a time seems to rapidly increase the region of similar slopes for the controllers’ steady state response curves. For this modified controller, integration leakiness is defined to incorporate the second annihilation rate constant *η*_2_

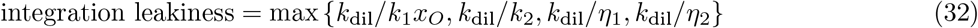

To systematically test whether the modified antithetic controller actually tolerates much higher integration leakiness than the original antithetic controller, we randomly generated 50 sets of controller parameters for both the original and the modified controller such that a controller steady state has at least 16 fold higher change compared to the a fixed fold change in system output *x*_*O*_. As predicted, the modified antithetic controller tolerated at least 10-fold higher dilution constant *k*_dil_ and at least 10-fold higher integration leakiness than the original antithetic controller. This higher integration leakiness tolerance vastly reduces the metabolic burden (*24*) required to maintain homeostatic performance. Although the previous published brink controller consisting of the antithetic controller and a downstream switch *x*_4_ (*7*)

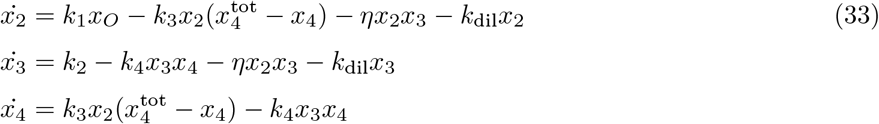

also showed increased tolerance toward the dilution constant, such tolerance completely disappeared when the downstream switch was also subject to the same dilution (Fig. 6F)

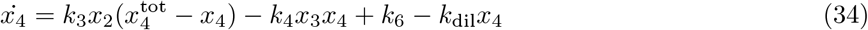

Sameniego et al (*7*) similarly reported that including dilution within the downstream switch *x*_4_ can degrade the brink controller’s near-homeostasis performance. In summary, inverse homeostasis plots allowed us to rationally modify the antithetic controller to truly increase the controller’s tolerance toward dilution, by increasing the range of slope overlap between steady state response curves of the two controller components toward each other.

The modified antithetic controller can be implemented using intein-gated dimer-based inactivation of the active domains (Fig. 6G). Each of the two controller proteins *x*_2_, *x*_3_ contains two orthogonal split inteins (*25*) on both the N and C terminus, its own partial monomer cargo, an activity monomer, and a cargo activator for the other controller’s cargo. The split inteins allow complete and irreversible monomer cargo activation at either N or C terminus. Forming low amount of active cargo only somewhat hinders the ability for two activity monomers to come together. However, binding between two active cargos of the same type bends the dimerized protein in a way prevent activity monomers from coming together, giving rise to the apparent cooperative elimination of protein activity 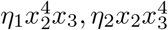.

## Discussion

The knowledge gained from finding a fast and reliable optimization method to find near-homeostasis supporting topologies as well as the novel methods to characterize near-homeostasis mechanisms both form a solid foundation for finding nearly-homeostatic systems with different transient response requirements. For instance, although the goal of an automated insulin delivery device is to maintain glucose steady state despite different internal environments within each patient, it is crucial for the insulin delivery device to avoid over-delivery of insulin that increases the chance of medically dangerous hypoglycemia. Insulin over-delivery arise from the delay between insulin injection and insulin action, as well as the lack of knowledge in a patient’s internal environments to accurately calculate the amount of insulin to inject. It would be useful to see which of the existing near-homeostasis mechanisms would be best at avoiding insulin over-delivery. At the same time, it would be wise to apply a modified version of our optimization approach to search for a nearly-homeostatic system that is good at avoiding insulin over-delivery.

In conclusion, out of the four formulations of the constrained optimization problem, we found two formulations that discovered near-homeostasis supporting architectures much faster and as comprehensively as brute-force search. For each discovered near-homeostasis supporting parameter set for both large genetic regulatory networks and large chemical reaction networks, we then utilized two levels of inverse homeostasis plots to analyze how a very small fold change in steady state system output is ultimately induced by a relatively large fold change in perturbation. Such analysis allowed us to identify mechanisms underlying the vast majority of near-homeostasis supporting parameter sets. We identified a novel two-component crossactivation controller with autocatalysis that tolerates much higher undesired basal expression levels than the well-known single-component autocatalytic controller. Furthermore, analysis of the antithetic controller using inverse homeostasis plots inspired further modifications to the antithetic controller that truly increased its tolerance toward undesired first order dilution rates. Our comprehensive toolset for the discovery and analysis of near-homeostasis-supporting architectures would be valuable for designing cellular therapeutics that maintain consistent performance in different patient environments.

## Materials and methods

### Abbreviations and definitions

**Antithetic integral controller** Eqn 23

**Antithetic integral controller with dilution** Eqn 25

**Arbitrary network and its derivatives** 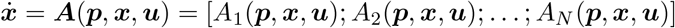, where ***p*** ∈ *P* ⊆ ℝ^dim(***p***)^, ***x*** ∈ *X* ⊆ ℝ^*N*^, ***u*** ∈ *U* ⊆ ℝ^dim(*u*)^, and *x*_*O*_ = *h*(***p, x***) is the scalar output of the system.

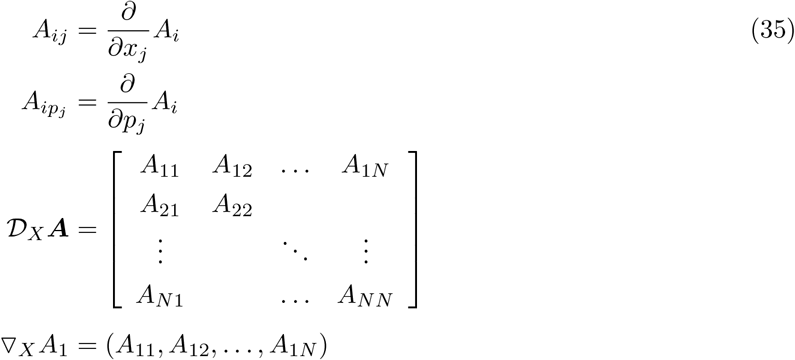

**Brink controller** Eqn 33

**Brink controller with dilution** Eqn 34

**Empirical relative sensitivity** Eqn 4

**Genetic network** Eqn 19

**Genetic network with continuous extension** Eqn 20

**Matrix indices convention** Suppose ***M*** is a 3 by 3 matrix of real numbers

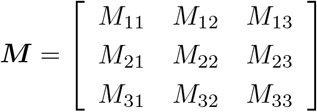

***M*** (*i, j*) refers to the entry in the i-th row starting from the top and j-th column starting from the left.

***M*** ([1, 2], [1, 2]) is the submatrix consisting of the first two rows and columns of ***M***.

**Modified antithetic controller** Eqn 31

**Network architecture** the combination of a topology and a class of interaction strengths (i.e. non-topological parameter sets) associated with that topology.

**Network topology** the set of qualitative interaction patterns (e.g. activation and repression) within a regulatory network.

**One component autocatalytic controller** Eqn 21

**Relative sensitivity** Eqn 3

**Reaction network rates** Fig. 2E, 2F

**Reaction network rates with continuous extensions** Eqn 11

**Parameter set** composed of a set of qualitative interaction types between different system components (defined as topological parameters) and a set of quantitative regulatory strengths for those interactions (defined as non-topological parameters).

**Scalar vs vector quantifies** The bolded ***x*** implies a vector quantity and the non-bolded *x* implies a scalar quantity.

**Two component cross-activation controller** Eqn 22

### The naive approach to find near-homeostasis supporting parameter sets

Starting from a random set of parameters ***p***_**0**_ and a fixed perturbation ***u***, we first estimate the steepest descent vector for the truncated relative sensitivity function using the finite differences. Suppose there are only two parameters (i.e. dim(***p***) = 2). We set *δ* > 0 to a very small number, and simulate the steady state

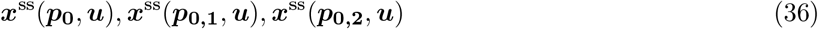

for each of the perturbed parameter sets ***p***_**0**_, ***p***_**0**,**1**_ = ***p***_**0**_ + [*δ*; 0], ***p***_**0**,**2**_ = ***p***_**0**_ + [0; *δ*]. A steady for (***p, u***) is reached if solution within the user-defined cutoff interval *t*_cutoff_ = [1 × 10^16^, 2 × 10^16^] results in sufficiently small fluctuations for all system components. Both the sensitivity function and the relative sensitivity function can be extended to be defined at all system component levels, not just the steady states

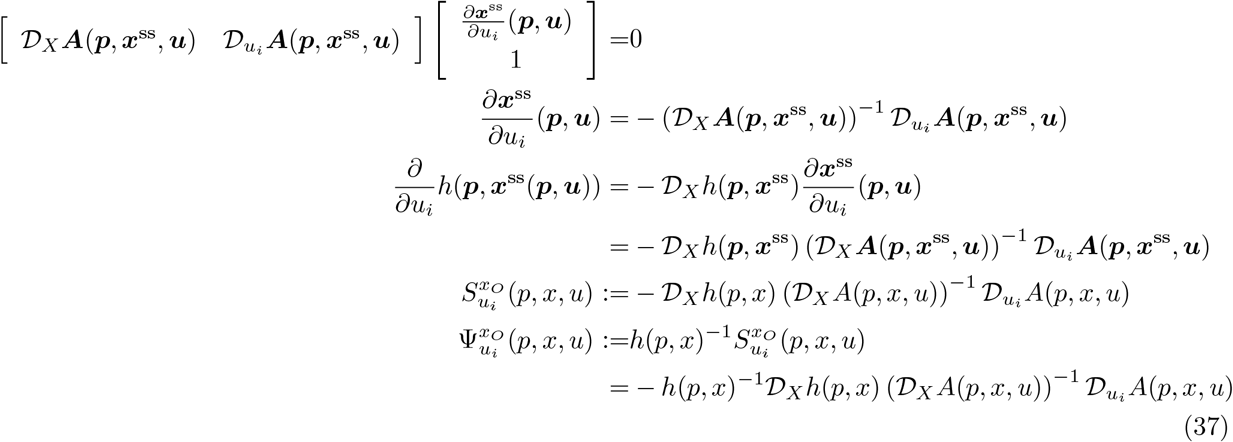

The total truncated relative sensitivity

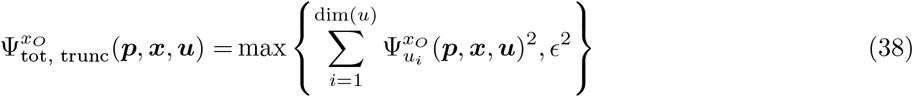

can be calculated for each of the perturbed parameter sets {***p***_**0**_, ***p***_**0**,**1**_, ***p***_**0**,**2**_} to result in 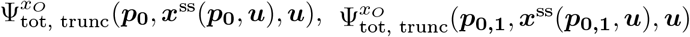, and 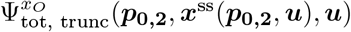. The parameter vector that results in the steepest change in the total truncated relative sensitivity is

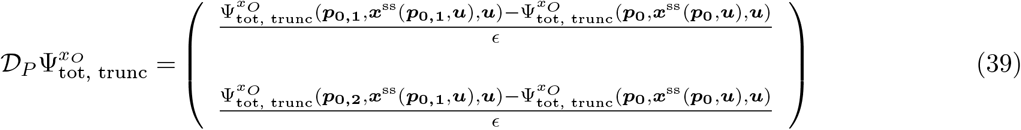

If a steady state cannot be reached for (***p, u***), then the truncated relative sensitivity is set to NaN.

Once the steep descent vector 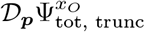 is found, a quasi-Newton step is taken to try to locate the parameter ***p*** such that the multi-dimensional slope of the objective function is zero

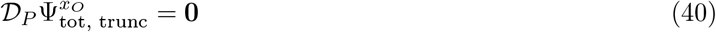

If steepest descent vector function 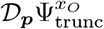 is linear and its multi-dimensional slope (Hessian) is easily computable, then desired parameter set ***p*** from the Newton step can be directly computed from the starting parameter set ***p***_**0**_

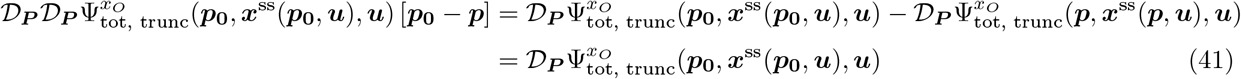

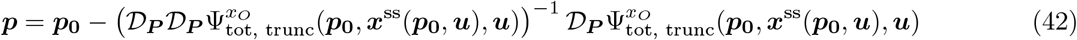

Since steepest descent vector function 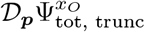 is often not linear, Newton steps are repeated until the multidimensional slope of the objective function 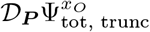 is sufficiently small. Newton steps are often enormous, giving rise to the seemingly erratic parameter trajectory for the often non-quadratic objective function 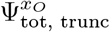 (Fig. 3G). Since estimating the Hessian using finite differences further involves dim(*p*)^2^ iterations of steepest descent vector estimation, the Hessian is estimated using the BFGS algorithm (*26*) within fmincon instead of using finite differences.

### The Lagrange method with rate vector magnitude constraint

In this method, we want to find the stationary point of the Lagrangian that uses the total truncated relative sensitivity function as the objective function and rate vector’s magnitude as the single nonlinear constraint

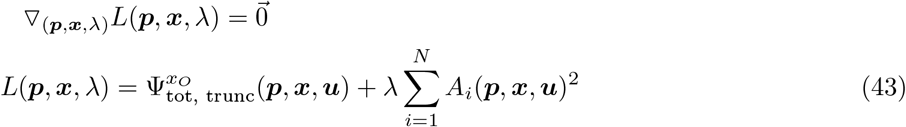

but analytical expression for the gradient of the total truncated relative sensitivity function is non-trivial to compute. The hardest part of computing the total truncated relative sensitivity function’s gradient is computing the gradients of the relative sensitivity function for an individual perturbation

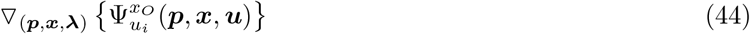

If *x*_1_ is the system output, the perturbation only affects *x*_1_’s rate, and the traditional expression of the relative sensitivity function using the adjugate matrix were used (see Eqn 37 for its derivation; (*1*, 2, 27, 12))

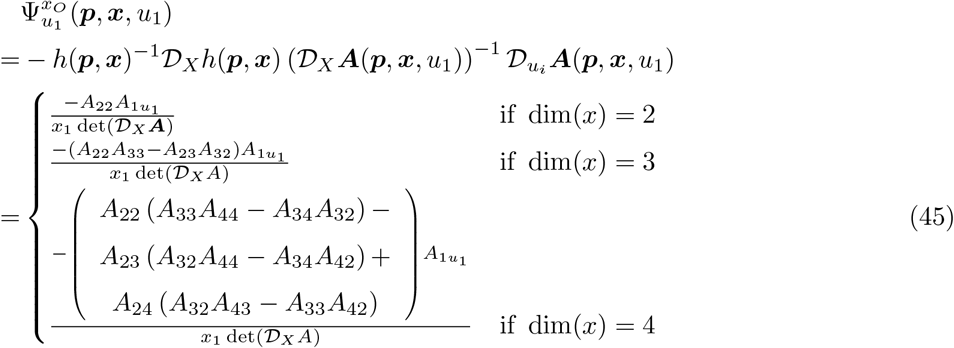

then the relative sensitivity function’s gradient cannot be computed in polynomial time due to the rapid increase in the number of additive terms.

Alternatively, we can implicitly compute a relative sensitivity function’s gradient. Let *ξ* be either a parameter or system component. The partial derivative of the relative sensitivity function involves the partial derivative of the sensitivity of every system component in response to all perturbations

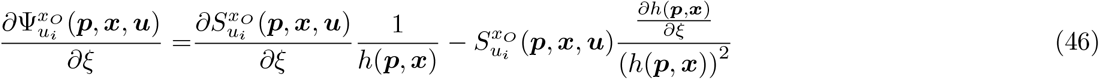

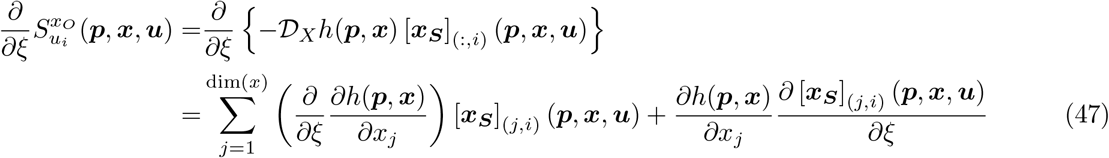

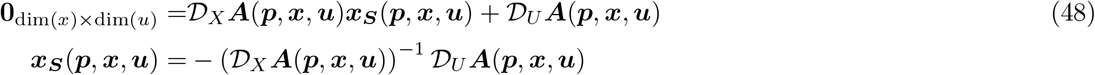

In turn, we can compute the sensitivity matrix’s partial derivative 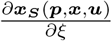 as the solution to a matrix equation by taking the derivative of the sensitivity matrix relation (Eqn 48)

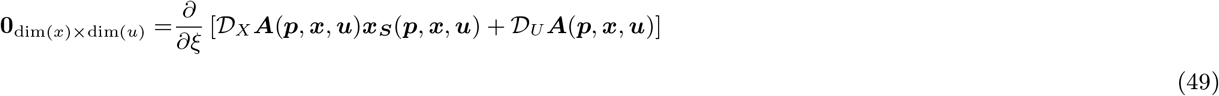

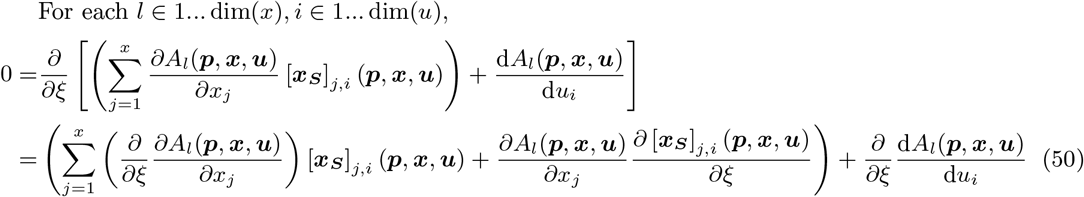

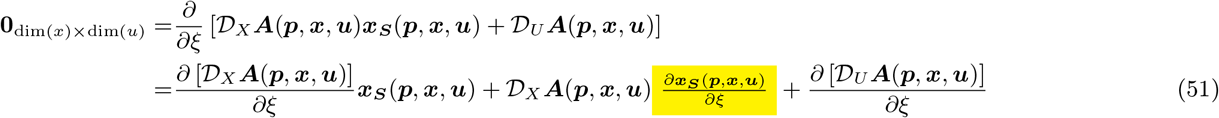

This seemingly more complex way to compute the relative sensitivity function’s partial derivatives is actually bounded by polynomial time, with asymptotic complexity of Θ(2 dim(***x***)^3^) after accounting for both the matrix multiplication and taking the matrix inverse of 𝒟_*X*_ ***A***(***p, x, u***).

### The gradient descent within constraints approach

The gradient descent within constraints approaches tries to find a near-homeostasis supporting parameter set by walking within the parameter-steady-state hyperspace down the steepest parameter direction that decreases the total truncated relative sensitivity

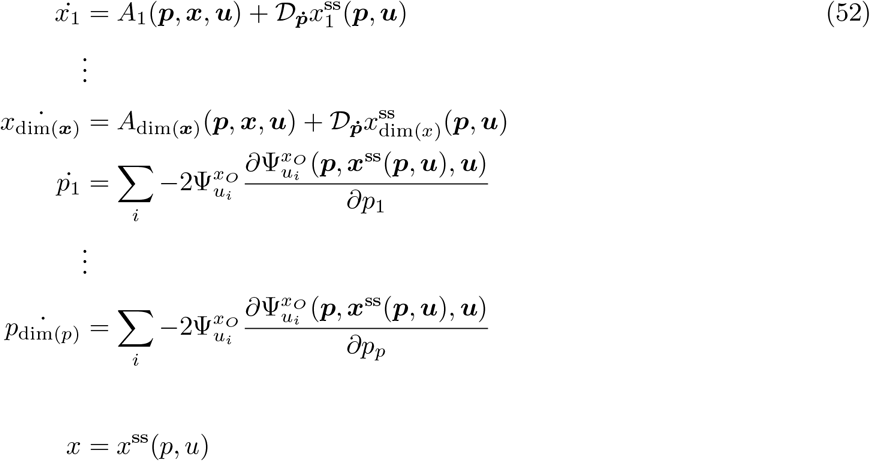

Here, even if the system component levels ***x*** is not at a steady state of the original system, the partial derivative of the relative sensitivity function with respect to parameters

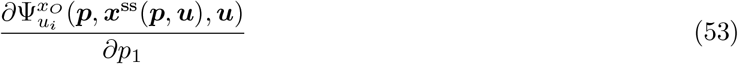

takes into account how the parameter change would change the system component levels as if they were at steady state. In contrast, the the corresponding partial derivative in Lagrange method with rate vector magnitude constraint approach

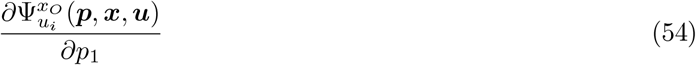

assumes that system component levels ***x*** does not change as the parameter *p*_1_ changes. The directional derivative

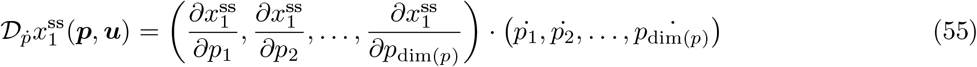

calculates how much the system component *x*_1_ is expected to change after parameters change. The original rates *A*_1_(***p, x, u***), …, *A*_dim(*x*)_(***p, x, u***) within the rate equations aims to steer the current state toward a steady state before any parameter change.

The partial derivative of the steady state relative sensitivity function (Eqn 53) is also computed implicitly like the Lagrange method with rate vector magnitude constraints, but additionally takes into account how the system component levels ***x*** would change in response to parameter change as if ***x*** is a steady state. The partial derivative of the steady state relative sensitivity function involves both the parameter partial derivative of steady state sensitivity function and steady state sensitivity functions in response to the chosen parameter *p*_1_

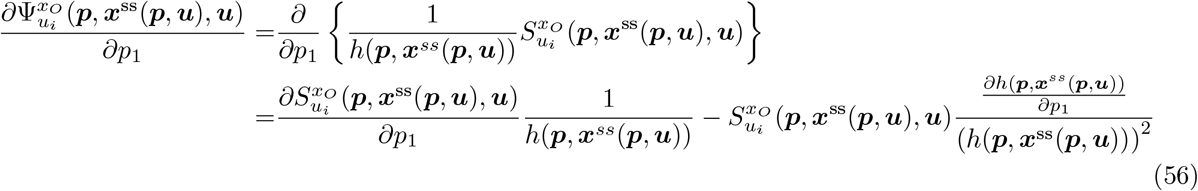

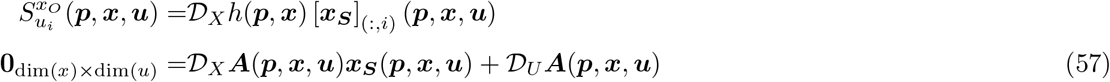

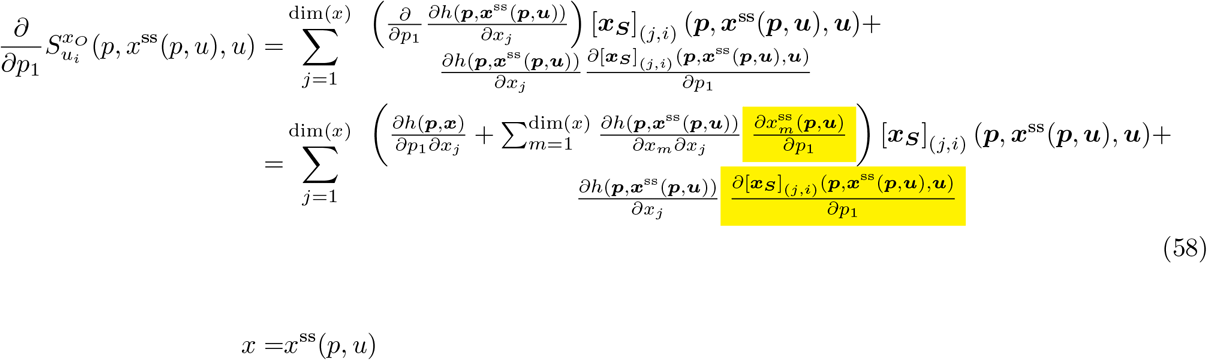

In turn, we can compute the sensitivity matrix’s partial derivative 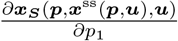 as the solution to a matrix equation by taking the derivative of the sensitivity matrix relation (Eqn 57).

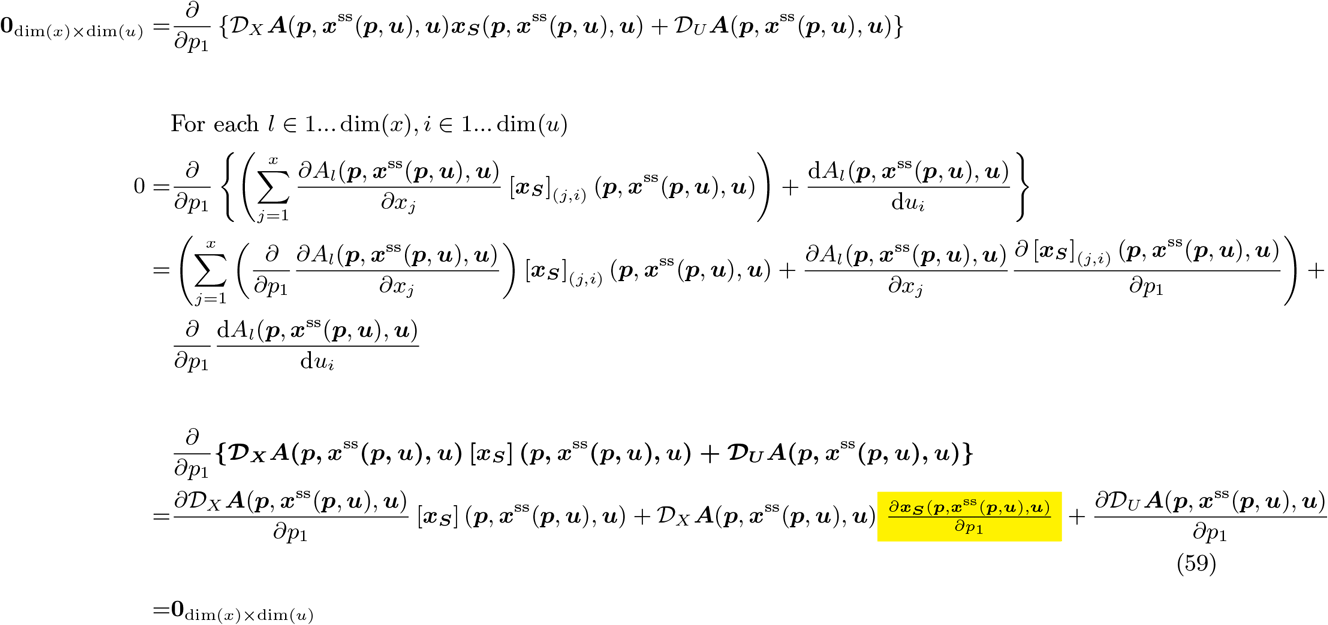

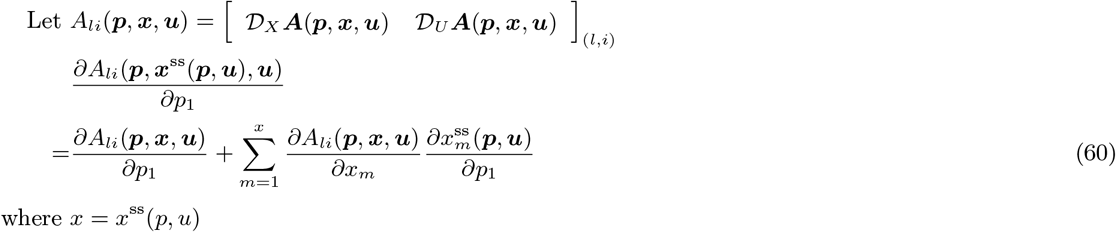

### The Lagrange method with rates constraints

Matlab’s fmincon is used to find the stations point of the Lagrangian that uses the total truncated relative sensitivity function as the objective function and all rates equations {*A*_1_(***p, x, u***), *A*_2_(***p, x, u***), …, *A*_*N*_ (***p, x, u***)} as multiple constraints

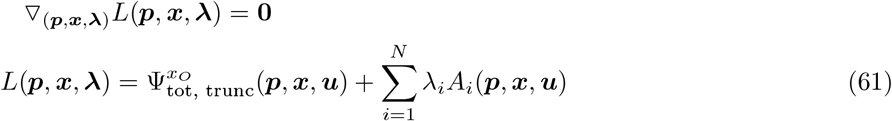

All gradients are computed in the same manner as the Lagrange method with rate magnitude constraints.

### Finding and analyzing near-homeostasis supporting parameter sets using a custom Matlab package

We built a software package BioSystem Suite in Matlab to perform simulation of a regulatory network and wrap all information related to each regulatory network into a single BioSystem or subclass (e.g. Genet-icNetwork, RNetwork) object. We will provide an illustrating example as part of our software overview.

Each rate equation for each system component can be specified using a symbolic expression within the “sys” property

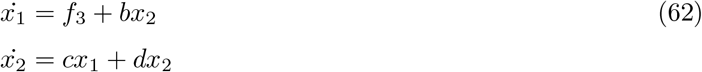

Rate equations may contain symbolic variables that are defined by symbolic expressions within the “flux” property. All symbolic expressions, including those within the “flux” property, can contain symbolic variables that are further defined by the “flux” property

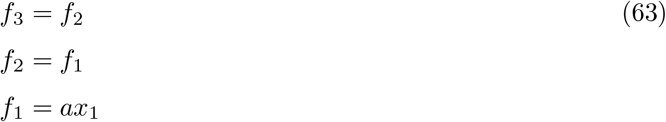

In addition, rate equations may contain symbolic variables whose numerical values can be specified in the “param” property {*a, b, c*}. Symbolic variables whose numerical values are not specified in the “param” property (*d*) are automatically determined to be a member of “inputvars” (also known as perturbations). The time-dependent perturbation values can be specified in the “ut” property. The output of the system can be specified by a symbolic expression within the “output_label” and “output” property

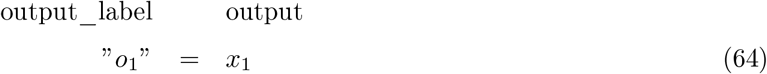

The range of allowable values for each system component is specified in the “xrange” property. The range of allowable values for each parameter is specified in the “prange” property. The initial conditions are specified in the “x0_map” property.

The software package allows rapid change in initial condition, system component ranges, parameter values, parameter value ranges, and time-dependent perturbation values by setting the “locked” property to true. To simulate system component levels across time, Matlab requires the system to be collapsed into a single set of equations when the property “num_sim_only” is false

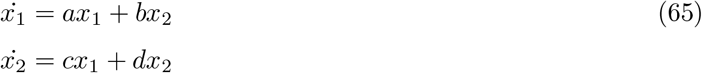

or the expressions arranged in sequentially executable manner

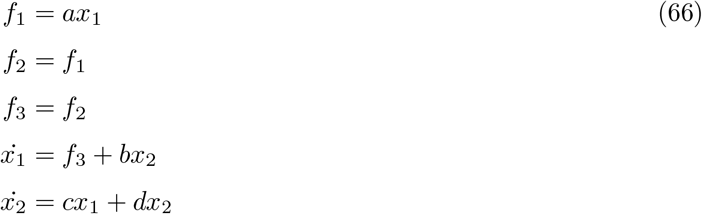

when the property “num_sim_only” is true. The numerically computable function used in simulations is stored in the “numeric_form” and “numeric_form_full” properties. The numerically computable output function is stored in the “numf_output” and “numf_output_full” properties. However, collapsing the system or arranging expressions for each simulation takes an unnecessarily long amount of time. We implemented the “locked” property that stores the collapsed system or arranged expressions to prevent recomputation and blocks attempts to change the system in a way that invalidates the stored numerical functions.

The software package contains convenient features to find near-homeostasis supporting parameter sets and analyze their underlying mechanisms of near-homeostasis, for arbitrary regulatory networks with differentiable rate equations. Features relevant for the purpose of the current study are:

- “convert_to_process” removes numeric symbolic variables from the “param” property such that those variables become perturbations.
- “get_feasible” randomly sets parameter values in the “param” property such that the supplied constraint is satisfied. In particular, supplying an always-true constraint allows one to completely randomize parameter values in the “param” property.
- “simulate_sys” determines the time-dependent system component values of the regulatory system.
- “find_steadystate” finds steady states of the regulatory system, and “find_dose_response” does the same but at different persistent perturbation levels.
- “gen_augsys4nearh_pxrsenso1min” generates an augmented object of the same BioSystem subclass that computes the gradient of the total truncated relative sensitivity function with respect to regulatory parameters and system components (Eqn 51). The “reverseAutoDiff” helper function is used to perform reverse automatic differentiation.
- “gen_augsys4nearh_prsenso1min_novs” generates an augmented object of the same BioSystem subclass that computes the gradient of the total truncated relative sensitivity function with respect to regulatory parameters. This augmented system computes how steady states should be changed based on change in parameter values (Eqn 59). The “reverseAutoDiff” helper function is used to perform reverse automatic differentiation.
- “findnhp_naive” finds near-homeostasis supporting continuous parameter sets using the naive approach outlined in the above sections. This algorithm uses the augmented system generated by “gen_augsys4nearh_pxrsenso1min”. “find_dose_response” is used to verify near-homeostasis of a candidate parameter set.
- “findnhp” finds near-homeostasis supporting parameter sets using the gradient descent within constrains approach outlined in the above sections. This algorithm uses the augmented system generated by “gen_augsys4nearh_prsenso1min_novs”. “find_dose_response” is used to verify near-homeostasis of a candidate parameter set.
- “findnhp_fmincon” finds near-homeostasis supporting parameter sets using either the Lagrangian with rate magnitude constraint method or the Lagrangian with rates constraints method. The Lagrangian with rate magnitude constraint method is selected when the mode argument contains “sqrt_sumsqsys constraint”. This algorithm uses the augmented system generated by “gen_augsys4nearh_pxrsenso1min”. “find_dose_response” is used to verify near-homeostasis of a candidate parameter set.
- “compute_invhomeo” computes level 1 inverse homeostasis plots based on the supplied steady state anchor points. Production and degradation rates of the system are required. For certain subclasses containing highly similar members (e.g. GeneticNetwork, RNetwork), the “gen_proddegrates” method can be used to automatically compute the correct production and degradation rates.
- “compute_invhomeo_lvl2” computes level 2 inverse homeostasis plots based on the supplied steady state anchor points.

One can see how the above features in combination with different simulation parameters are used to generate different figures in this study (Table 1). Additional example usages of those figures can be found in relevant tests in the BioSystemTests unit testing framework.

**Table 1:**
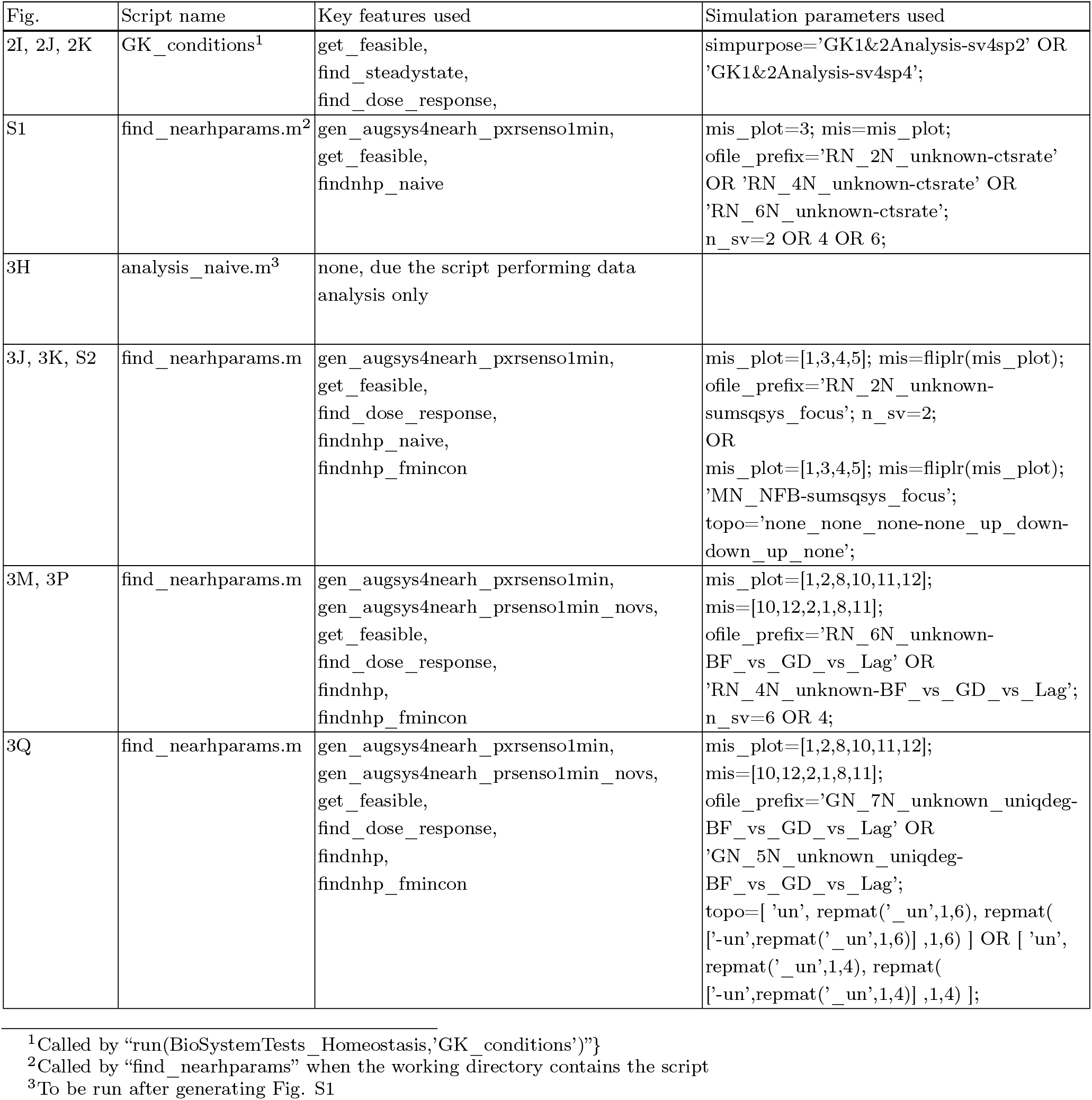
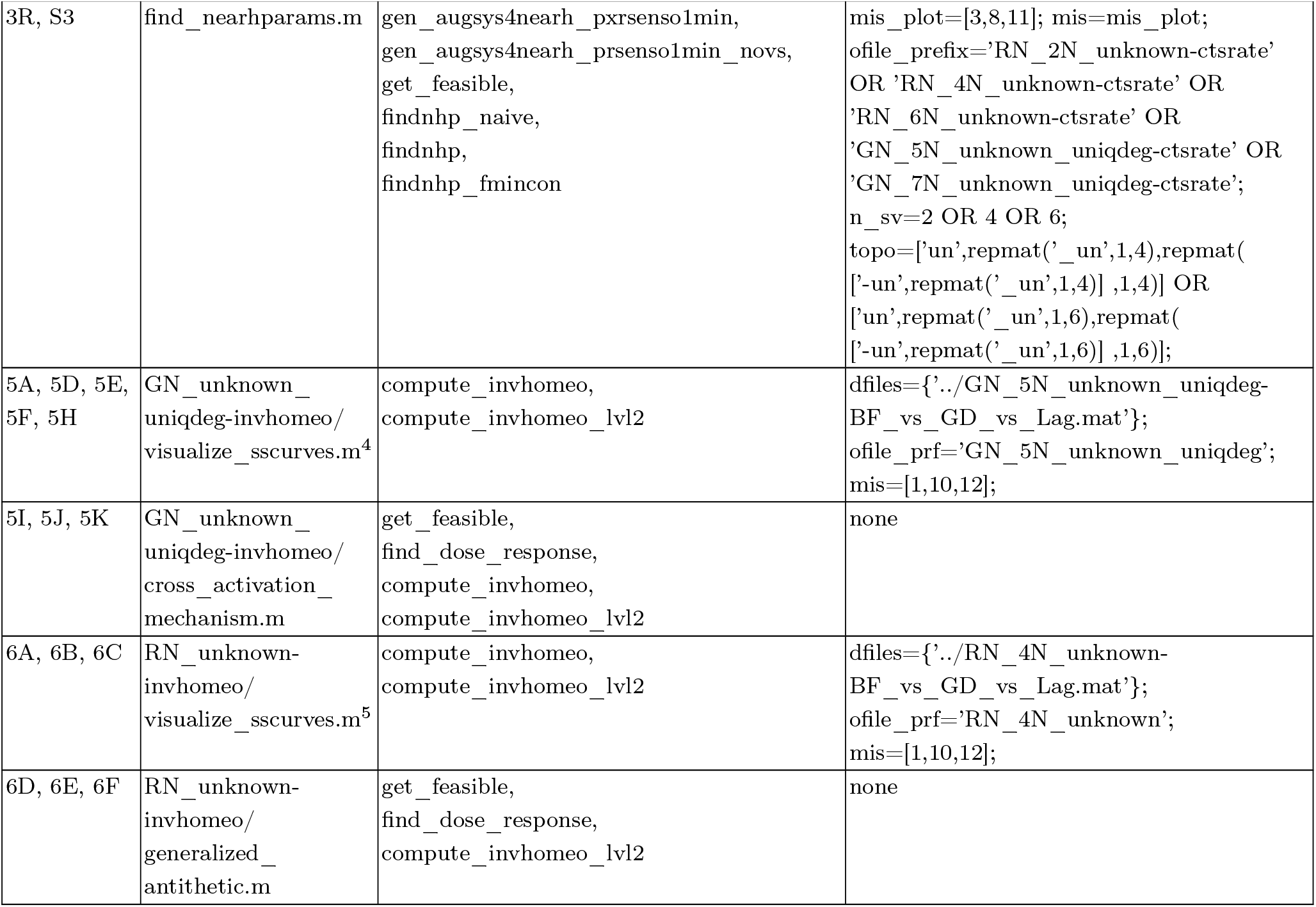
Simulation parameters used to generate different figures.

Each search iteration was analyzed to see whether the iteration achieved its intended goal and why the intended goal may not have been achieved. Different classes of search iterations are present in Table 2.

**Table 2:**
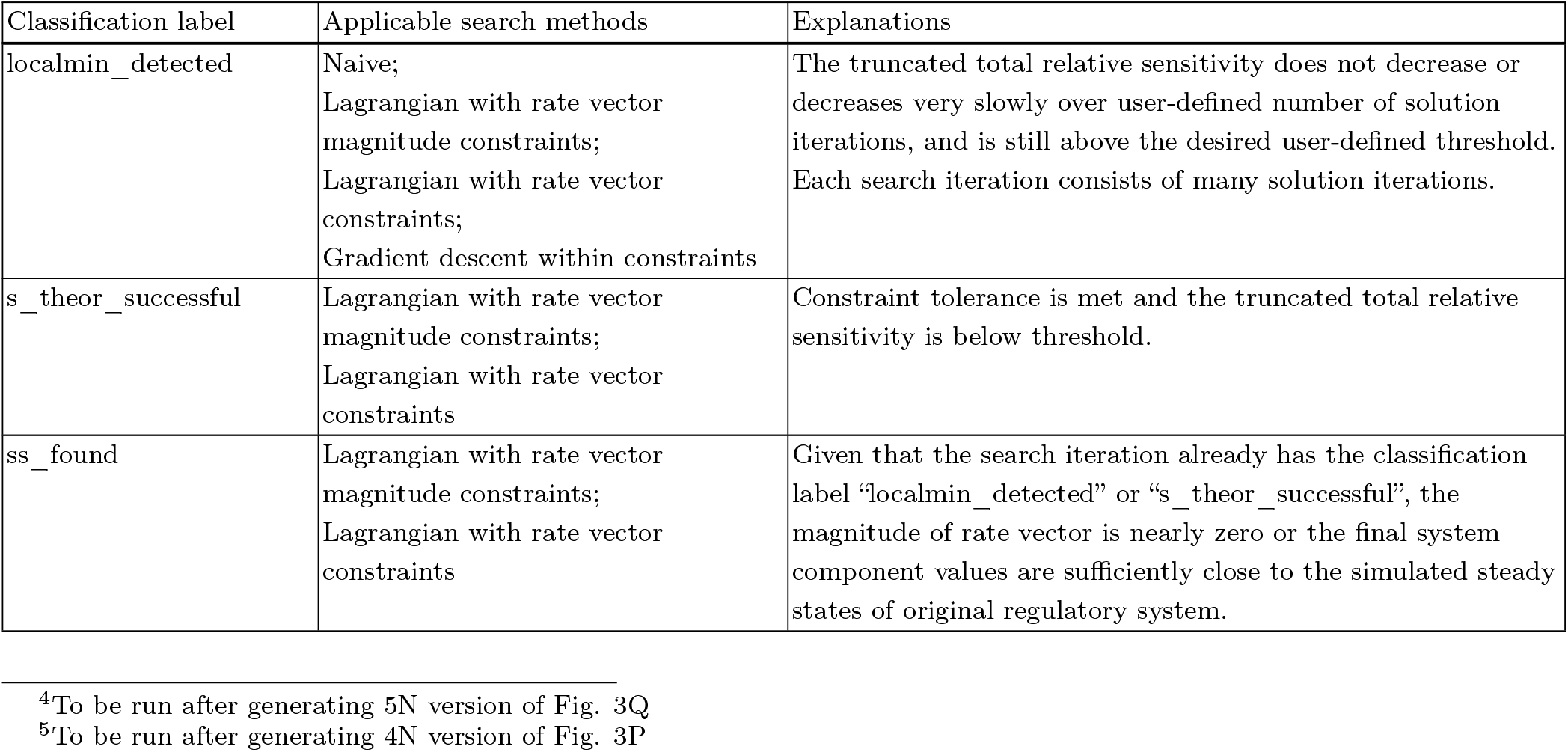
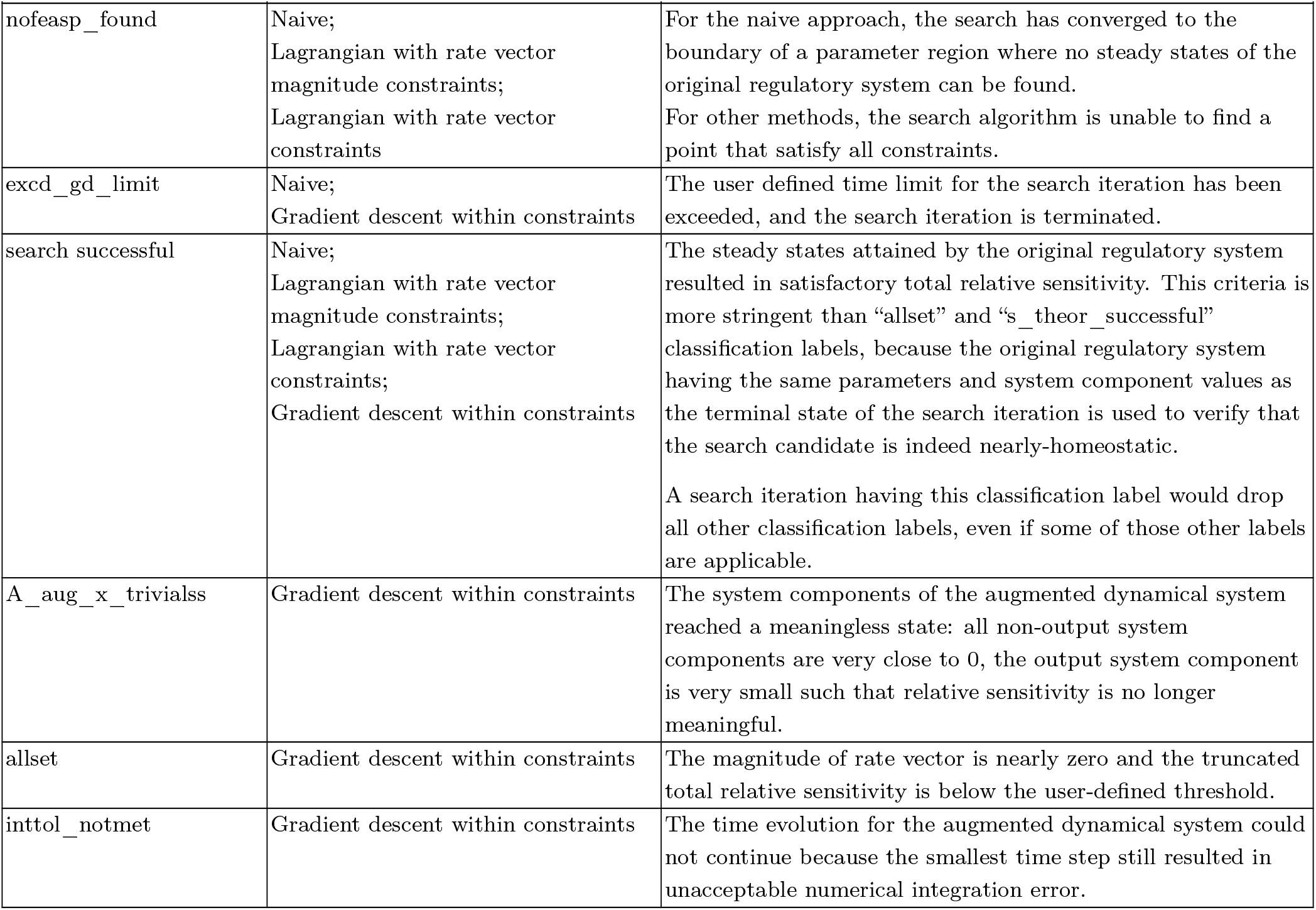
Explanations for different classes of search iterations.

## Author contributions

Z.T. conceived the presented ideas. Z.T. and D.R.M implemented and evaluated the ideas. Z.T. and D.R.M wrote the paper.

## Data and materials availability

All code used to generate the simulation data in the figures are contained in the SupplementaryCode zip folder.

## Supporting text

**Fig. S1:**
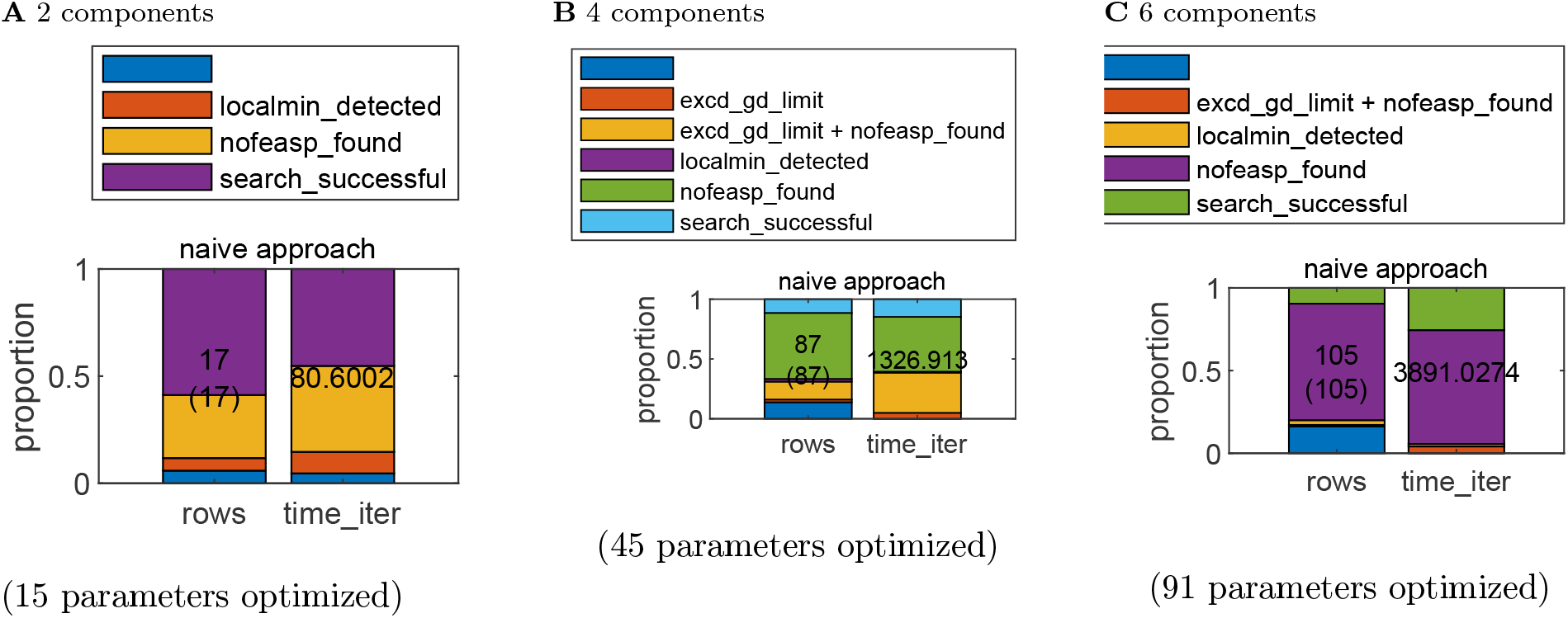
Detailed characterization of the naive approach to find near-homeostasis supporting parameter sets for continuously extended reaction networks. Please see Methods for a detailed explanation of classification labels.

**Fig. S2:**
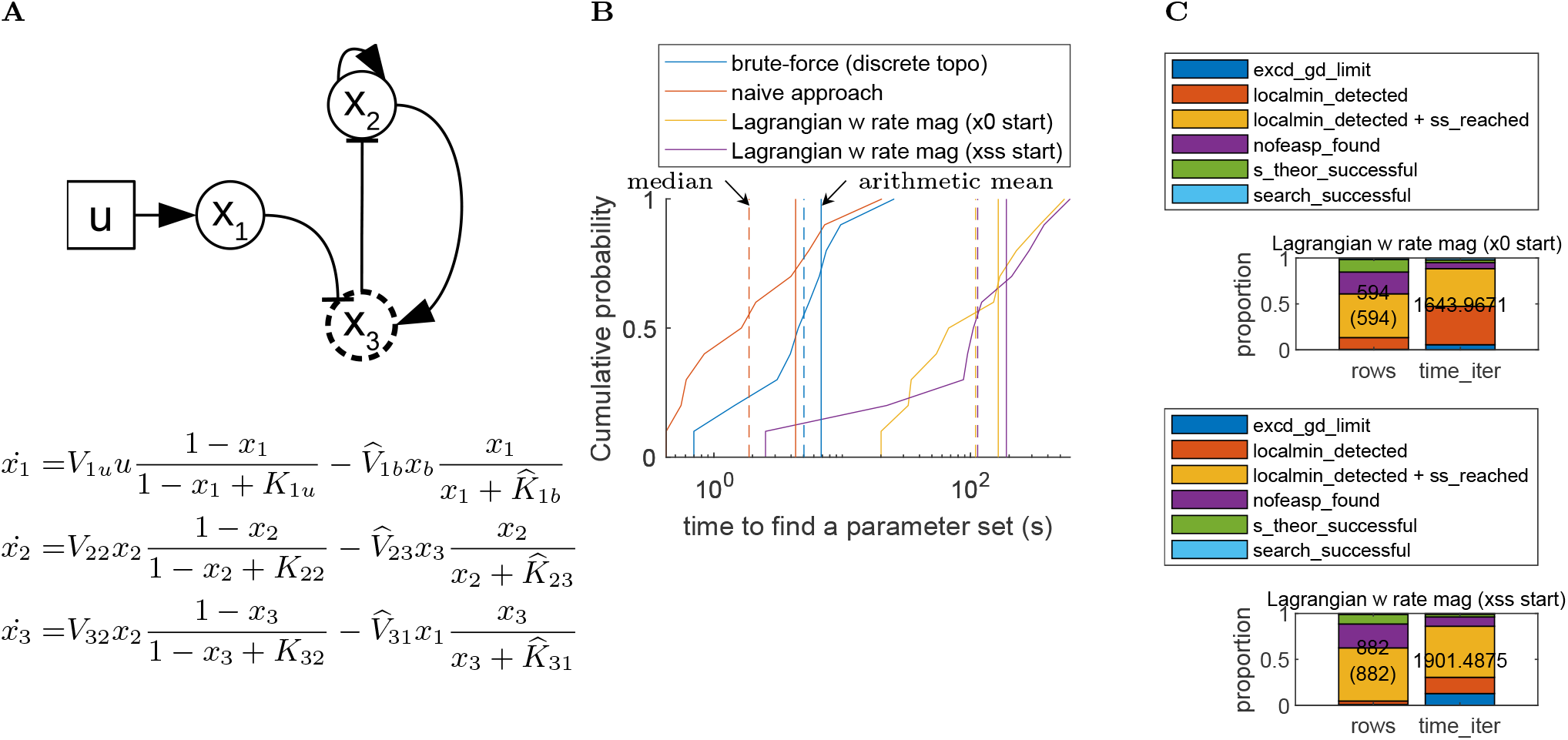
Applying the Lagrangian method with rate vector magnitude constraints to find near-homeostasis supporting parameter sets for Ma et al (*1*)’s negative feedback loop. Please see Methods for a detailed explanation of classification labels.

**Fig. S3:**
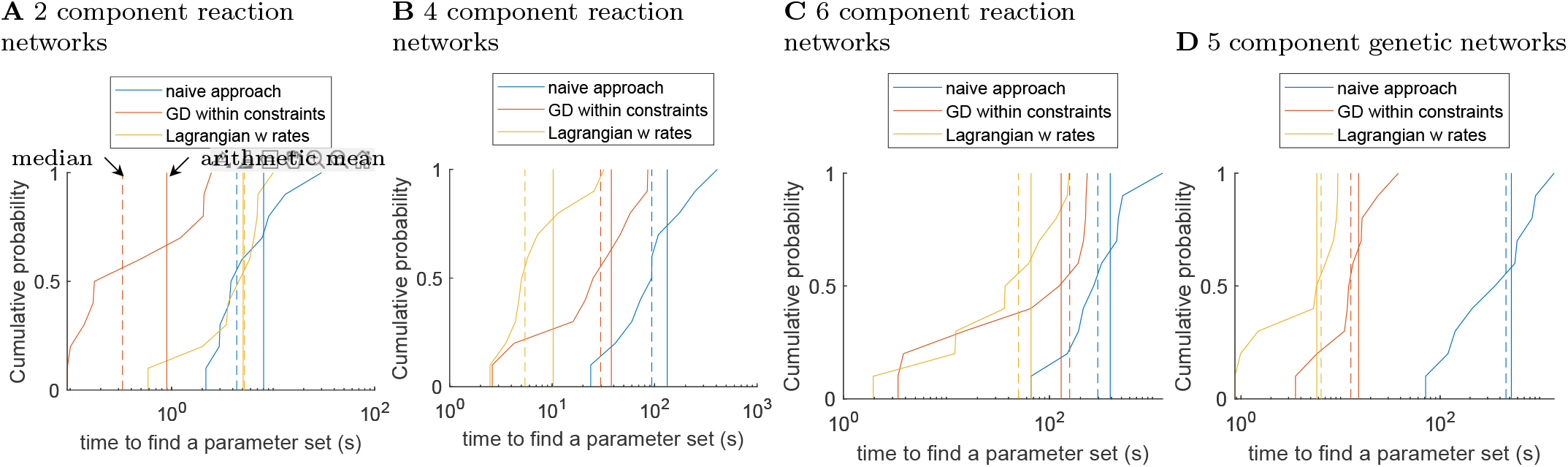
Comparing the naive approach, gradient descent within constraints, and the Lagrange method with rates constraints in terms of how fast a near-homeostasis continuous parameter set can be found.

## References

1. W. Ma, A. Trusina, H. El-Samad, W. A. Lim, C. Tang, Defining Network Topologies that Can Achieve Biochemical Adaptation. Cell 138 (4), 760–773 (2009), doi:10.1016/j.cell.2009.06.013, http://www.cell.com/abstract/S0092-8674(09)00712-0.

2. Z. F. Tang, D. R. McMillen, Design principles for the analysis and construction of robustly homeostatic biological networks. Journal of Theoretical Biology 408, 274–289 (2016), doi:10.1016/j.jtbi.2016.06.036.

3. T. Peyser, E. Dassau, M. Breton, J. S. Skyler, The artificial pancreas: current status and future prospects in the management of diabetes. Annals of the New York Academy of Sciences 1311, 102–123 (2014), doi:10.1111/nyas.12431.

4. M. Breton, et al., Fully Integrated Artificial Pancreas in Type 1 Diabetes. Diabetes 61 (9), 2230–2237 (2012), doi:10.2337/db11-1445, https://www.ncbi.nlm.nih.gov/pmc/articles/PMC3425406/.

5. W. Shi, W. Ma, L. Xiong, M. Zhang, C. Tang, Adaptation with transcriptional regulation. Scientific Reports 7, srep42648 (2017), doi:10.1038/srep42648, https://www.nature.com/articles/srep42648.

6. C. Briat, A. Gupta, M. Khammash, Antithetic Integral Feedback Ensures Robust Perfect Adaptation in Noisy Biomolecular Networks. Cell Systems 2 (1), 15–26 (2016), doi:10.1016/j.cels.2016.01.004, http://www.cell.com/article/S2405471216000053/abstract.

7. C. C. Samaniego, E. Franco, Ultrasensitive molecular controllers for quasi-integral feedback. Cell Systems (2021), doi:10.1016/j.cels.2021.01.001, https://www.sciencedirect.com/science/article/pii/S2405471221000351.

8. E. D. Sontag, Adaptation and regulation with signal detection implies internal model. Systems & Control Letters 50 (2), 119–126 (2003), doi:10.1016/S0167-6911(03)00136-1, https://www.sciencedirect.com/science/article/pii/S0167691103001361.

9. A. Gupta, M. Khammash, Universal structural requirements for maximal robust perfect adaptation in biomolecular networks. Proceedings of the National Academy of Sciences of the United States of America 119 (43), e2207802119 (2022), doi:10.1073/pnas.2207802119.

10. R. P. Araujo, L. A. Liotta, Universal structures for adaptation in biochemical reaction networks. Nature Communications 14 (1), 2251 (2023), number: 1 Publisher: Nature Publishing Group, doi:10.1038/s41467-023-38011-9, https://www.nature.com/articles/s41467-023-38011-9.

11. R. Scheepers, N. L. Levi, R. P. Araujo, A distributed integral control mechanism for regulation of cholesterol concentration in the human retina. Royal Society Open Science 11 (10), 240432 (2024), doi:10.1098/rsos.240432.

12. Y. Wang, Z. Huang, F. Antoneli, M. Golubitsky, The structure of infinitesimal homeostasis in input-output networks. Journal of Mathematical Biology 82 (7), 62 (2021), doi:10.1007/s00285-021-01614-1.

13. J. Ang, S. Bagh, B. P. Ingalls, D. R. McMillen, Considerations for using integral feedback control to construct a perfectly adapting synthetic gene network. Journal of theoretical biology 266 (4), 723–738 (2010), doi:10.1016/j.jtbi.2010.07.034.

14. J. Ang, D. R. McMillen, Physical constraints on biological integral control design for homeostasis and sensory adaptation. Biophysical journal 104 (2), 505–515 (2013), doi:10.1016/j.bpj.2012.12.015.

15. T. Achterberg, R. Wunderling, Mixed Integer Programming: Analyzing 12 Years of Progress, in Facets of Combinatorial Optimization: Festschrift for Martin Grötschel, M. Jünger, G. Reinelt, Eds. (Springer, Berlin, Heidelberg), pp. 449–481 (2013), doi:10.1007/978-3-642-38189-8_18, https://doi.org/10.1007/978-3-642-38189-8_18.

16. OpenAI, et al., GPT-4 Technical Report (2024), doi:10.48550/arXiv.2303.08774, http://arxiv.org/abs/2303.08774, 2303.08774 [cs].

17. B. P. Ingalls, Mathematical Modeling in Systems Biology: An Introduction (The MIT Press, Cambridge, Massachusetts), 1 edition ed. (2013).

18. M. B. Elowitz, S. Leibler, A synthetic oscillatory network of transcriptional regulators. Nature 403 (6767), 335–338 (2000), doi:10.1038/35002125.

19. G. Lillacci, M. Khammash, Parameter Estimation and Model Selection in Computational Biology. PLOS Computational Biology 6 (3), e1000696 (2010), publisher: Public Library of Science, doi:10.1371/journal.pcbi.1000696, https://journals.plos.org/ploscompbiol/article?id=10.1371/journal.pcbi.1000696.

20. T. Drengstig, X. Y. Ni, K. Thorsen, I. W. Jolma, P. Ruoff, Robust adaptation and homeostasis by autocatalysis. The journal of physical chemistry. B 116 (18), 5355–5363 (2012), doi:10.1021/jp3004568.

21. S. K. Aoki, et al., A universal biomolecular integral feedback controller for robust perfect adaptation. Nature 570 (7762), 533 (2019), doi:10.1038/s41586-019-1321-1, https://www.nature.com/articles/s41586-019-1321-1.

22. H.-H. Huang, Y. Qian, D. D. Vecchio, A quasi-integral controller for adaptation of genetic modules to variable ribosome demand. Nature Communications 9 (1), 5415 (2018), doi:10.1038/s41467-018-07899-z, https://www.nature.com/articles/s41467-018-07899-z.

23. Y. Qian, D. Del Vecchio, Realizing ‘integral control’ in living cells: how to overcome leaky integration due to dilution? Journal of The Royal Society Interface 15 (139), 20170902 (2018), publisher: Royal Society, doi:10.1098/rsif.2017.0902, https://royalsocietypublishing.org/doi/10.1098/rsif.2017.0902.

24. B. Kell, R. Ripsman, A. Hilfinger, Noise properties of adaptation-conferring biochemical control modules. Proceedings of the National Academy of Sciences of the United States of America 120 (38), e2302016120 (2023), doi:10.1073/pnas.2302016120.

25. F. Pinto, E. L. Thornton, B. Wang, An expanded library of orthogonal split inteins enables modular multi-peptide assemblies. Nature Communications 11 (1), 1529 (2020), publisher: Nature Publishing Group, doi:10.1038/s41467-020-15272-2, https://www.nature.com/articles/s41467-020-15272-2.

26. R. R. Fletcher, Practical methods of optimization (Chichester ; New York : Wiley) (1987), http://archive.org/details/practicalmethods0000flet.

27. R. P. Araujo, L. A. Liotta, The topological requirements for robust perfect adaptation in networks of any size. Nature Communications 9 (2018), doi:10.1038/s41467-018-04151-6, https://www.ncbi.nlm.nih.gov/pmc/articles/PMC5931626/.

